# NeuWalk: A Python Library for Data-Driven Synthesis of Neuron Morphologies Using Random Walks

**DOI:** 10.64898/2026.09.25.754566

**Authors:** Francesco Cavarretta

## Abstract

Large-scale biophysically-detailed models of the brain are instrumental in unraveling how cellular, synaptic, and circuit-level mechanisms contribute to neural dynamics. Such models are built around large populations of single-neuron models whose morphologies must preserve physiological variability to reproduce integrative properties and firing patterns observed experimentally. Experimental reconstruction of morphologies is a time-consuming and resource-intensive process, unlikely to supply them at the required scale. Several data-driven methods for morphology synthesis have been developed, which nonetheless rely on extensive datasets for parameter estimation. To address this limitation, we developed NeuWalk, to our knowledge the first public Python library for morphology synthesis based on branching-and-annihilating, biased-and-correlated random walks. It estimates branching and annihilation rates from Sholl intersections and, optionally, bifurcation counts, while also ensuring their replication in synthetic morphologies; user-specified combinations of biases, implementing somatofugality, self-avoidance, spatial competition, and branching-angle control, sculpt morphological patterns characteristic of each neuron class. We then validated the method on mitral and middle tufted cells of the olfactory bulb, semilunar and pyramidal cells of the anterior piriform cortex, and neocortical pyramidal neurons: means and variances of Sholl intersections, bifurcation counts, and total dendritic length were statistically indistinguishable between synthetic and natural morphologies, with the single exception of its variance in apical dendrites of piriform cortex pyramidal neurons. As Sholl intersections and bifurcation counts are commonly reported in the literature, and biases can be inferred from published images or qualitative descriptions, NeuWalk requires no immediate access to experimental reconstructions, striking a trade-off between data-driven parameterization and empiricism without sacrificing morphological accuracy.

## 1. Introduction

Large-scale biophysically-detailed models of the brain are instrumental in investigating its functions by unraveling how cellular, synaptic, and circuit-level mechanisms contribute to neural dynamics [1, 2, 3, 4, 5, 6]. By integrating anatomical and electrophysiological constraints within a unified computational framework, these models enable simulation of brain activity with high realism. Also, their analysis can yield mechanistic explanations for the pathophysiology of brain diseases, providing the basis for testing specific hypotheses or innovative therapeutic strategies through simulations [7, 8, 9, 10].

Morphologically and electrophysiologically detailed single-neuron models constitute the backbone of large-scale biophysically-detailed brain models, which typically comprise hundreds or even thousands of neurons [1, 2, 3, 4, 5, 6]. Multiple studies have highlighted the influence of morphology on electrotonic and integrative properties, including cable filtering, synaptic integration, dendritic action potential propagation, and firing patterns [11, 12, 13, 14, 15, 16, 17, 18], together with its effects on neural dynamics [1, 3, 19, 20, 21, 22]. Additionally, morphological variability contributes to the diversity in firing patterns, information processing, and morpho-electric phenotypes [23, 24], while conferring robustness and adaptability to brain circuits [25, 26]. Hence, accurate simulation of neural dynamics requires the construction of these models around realistic morphologies that reproduce physiological variability. However, experimental reconstruction of morphologies is a time-consuming and resource-intensive process, requiring high-quality imaging, tracing of dendritic and axonal processes, correction of discontinuities, and extensive proofreading, and is therefore unlikely to supply morphologies at the scale required for large-scale brain modeling. To address this issue, several data-driven methods have been developed for morphology synthesis [27, 28, 29, 30]. However, most of these methods estimate parameters from sizable sets of experimental reconstructions, subordinating their usability to the availability of such datasets.

In previous publications, we used branching-and-annihilating, biased-and-correlated three-dimensional random walks to synthesize middle tufted and mitral cells [1, 3], two main populations of principal neurons in the olfactory bulb. Although the method produced realistic morphologies, it lacked mathematical rigor and required experimental reconstructions to estimate branching (i.e., bifurcation) and annihilation (i.e., branch termination) rates. To address this issue, we recently derived closed-form mathematical equations that allow the estimation of these rates from Sholl intersections (i.e., the relationship between the number of dendritic branches and the distance from the soma) and bifurcation counts (as an optional constraint) [31], which also constitute a method to replicate both measures in the synthesis of morphology based on random walks. Given how commonly Sholl intersections and bifurcation counts are reported in the literature, these equations support data-driven synthesis of morphologies without requiring direct access to reconstructions. However, this approach tended to yield higher variance than observed experimentally for both measures [31]. To correct this issue, we developed a bin-wise procedure that limits variance in synthetic morphologies, along with a calibration procedure that corrects the distortion thereby introduced in mean values. Here, we bring these ideas into a new software tool: *NeuWalk*, a public Python library for morphology synthesis based on branching-and-annihilating, biased-and-correlated random walks. Specifically, NeuWalk uses bifurcation and annihilation rates to synthesize sequences of elongation, bifurcation, and annihilation events (i.e., topology). At the same time, user-specified combinations of biases shape morphology by implementing somatofugality (i.e., outward orientation away from the soma), self-avoidance, and spatial competition, while correlation between successive step directions, along with randomness, sets trajectory smoothness. Notably, biases can be inferred empirically through visual inspection of published morphological images or qualitative descriptions.

Finally, we validated our method on mitral and middle tufted cells of the olfactory bulb, semilunar and pyramidal cells of the anterior piriform cortex, and neocortical pyramidal neurons. For each class, we compared Sholl intersections, bifurcation counts, and total dendritic length between synthetic and natural morphologies, and confirmed that they were statistically indistinguishable, with the single exception of the variance of total dendritic length in apical dendrites of piriform cortex pyramidal neurons.

In summary, we provide a data-driven method for morphology synthesis that tolerates missing experimental information (e.g., unavailability of morphological reconstructions or lack of estimates for bifurcation counts) while holding a reasonable degree of empiricism (e.g., biases regulating trajectories). The main contributions of this paper are:

- NeuWalk, to our knowledge the first public Python library for neuron morphology synthesis via branching-and-annihilating, biased-and-correlated random walks.
- Extension of the closed-form framework for the estimation of branching and annihilation rates from mono-dimensional validation to full threedimensional morphology synthesis.
- A progressive, bin-wise procedure with backtracking that corrects the variance excess observed in our previous formulation [31].
- Introduction of directional correlation as a mechanism for trajectory smoothness, alongside biases such as self-avoidance, spatial competition, and branching-angle control to shape morphological patterns characteristic of different neuron classes.
- Validation of full synthetic three-dimensional morphologies — including total dendritic length, not previously considered — against experimental reconstructions across different neuron classes.

We also emphasize that our effort does not aim to simulate biological processes underlying dendritic development, but rather to synthesize morphologies that are indistinguishable from natural counterparts.

## 2. Related work

In this section, we review several approaches for morphology synthesis developed in recent decades. L-Neuron applies recursive rules, drawing branch length, taper, diameter, branching angle, and bifurcation probability from experimental distributions, so that global features, such as the Sholl profile, emerge from local sampling [27]. NETMORPH models neurite outgrowth phenomenologically, with growth cones that elongate and branch stochastically according to rates calibrated against experimental data, and assembles the resulting arbors into networks [28]. In both cases, the estimation of distributions or rates required access to experimental reconstructions.

The Topological Neuron Synthesis method constrains bifurcation positions using the Topological Morphology Descriptor, which encodes a dendritic tree as the set of intervals spanned by dendritic branches, expressed as distances from the soma; the descriptor is extracted from experimental reconstructions [30]. The TREES Toolbox synthesizes morphologies by connecting *carrier points* through a custom version of the minimum spanning tree algorithm, under the assumption that these points represent putative synaptic targets [29, 32]. In this method, morphologies are shaped by a *balancing factor* that determines the trade-off between cable cost and conduction time [29, 32], which are proportional to total dendritic length and path-distance from the soma, respectively. Furthermore, the number of bifurcations correlates directly with the number of carrier points [32], which therefore constitutes an additional parameter. Both the balancing factor and the number of carrier points are determined empirically. Therefore, both methods rely on experimental morphological reconstructions, either to extract the descriptor or to tune parameters.

Recently, approaches relying on machine learning have emerged as an alternative paradigm. MorphVAE was the first such method, representing the path from soma to tip as a three-dimensional walk, synthesizing new walks with a sequence-to-sequence variational autoencoder, and assembling them into a full morphology by post-hoc clustering [33]. However, this clustering produces nodes with implausibly high branching degree, thereby invalidating part of the resulting morphologies, while branch angles and arbor extent are systematically underestimated [33]. MorphGrower addresses the issue by synthesizing morphologies layer by layer, synthesizing sibling branch pairs jointly and conditioning each new layer on the already-synthesized subtree, which enforces topological validity by construction [34]. However, this synthesis proceeds autoregressively with respect to a reference morphology that is required not only for training but also at synthesis time [34]. Therefore, both methods require access to experimental morphological reconstructions to train their models, and MorphGrower additionally requires a reference reconstruction at synthesis time.

An evolutionary-developmental approach synthesizes morphologies by simulating a developmental process, controlled by an artificial gene regulatory network [35]. In this method, morphology is represented as a sequence of interconnected cylindrical segments, in which each update of the network state corresponds to an elongation, bifurcation, or termination event of the growing segment. Network evolution is a multi-objective optimization, in which objectives quantify the differences between emergent morphological features of the virtual neurons and those measured from experimental reconstructions. Therefore, this method requires access to experimental morphological reconstructions for estimating the targets of the multi-objective function.

In summary, we have reviewed several approaches for morphology synthesis, noting that they depend on the availability of experimental reconstructions to extract parameters. None of them allows us to directly constrain synthesis to target morphometric statistics, such as Sholl intersections or bifurcation counts. Additionally, these methods are not fully automatic, since they require human supervision. In contrast, NeuWalk treats commonly reported morphometric statistics as constraints rather than emergent properties. It estimates bifurcation and annihilation rates from Sholl intersections, with bifurcation counts as an optional constraint, and ensures that both are reproduced by synthetic morphologies.

## 3. Methods

NeuWalk organizes morphology synthesis into multiple stages, illustrated in Figure 1. It begins by estimating Sholl profiles and bifurcation counts from experimental reconstructions of neuron morphologies (stage 1; see Section 3.1). These statistics yield bifurcation (β) and annihilation (α) rates for branching-and-annihilating mono-dimensional random walks (stage 2; see Section 3.2), which in turn synthesize sequences of elongation, bifurcation, and annihilation events (i.e., topology; stage 3).

**Figure 1:**
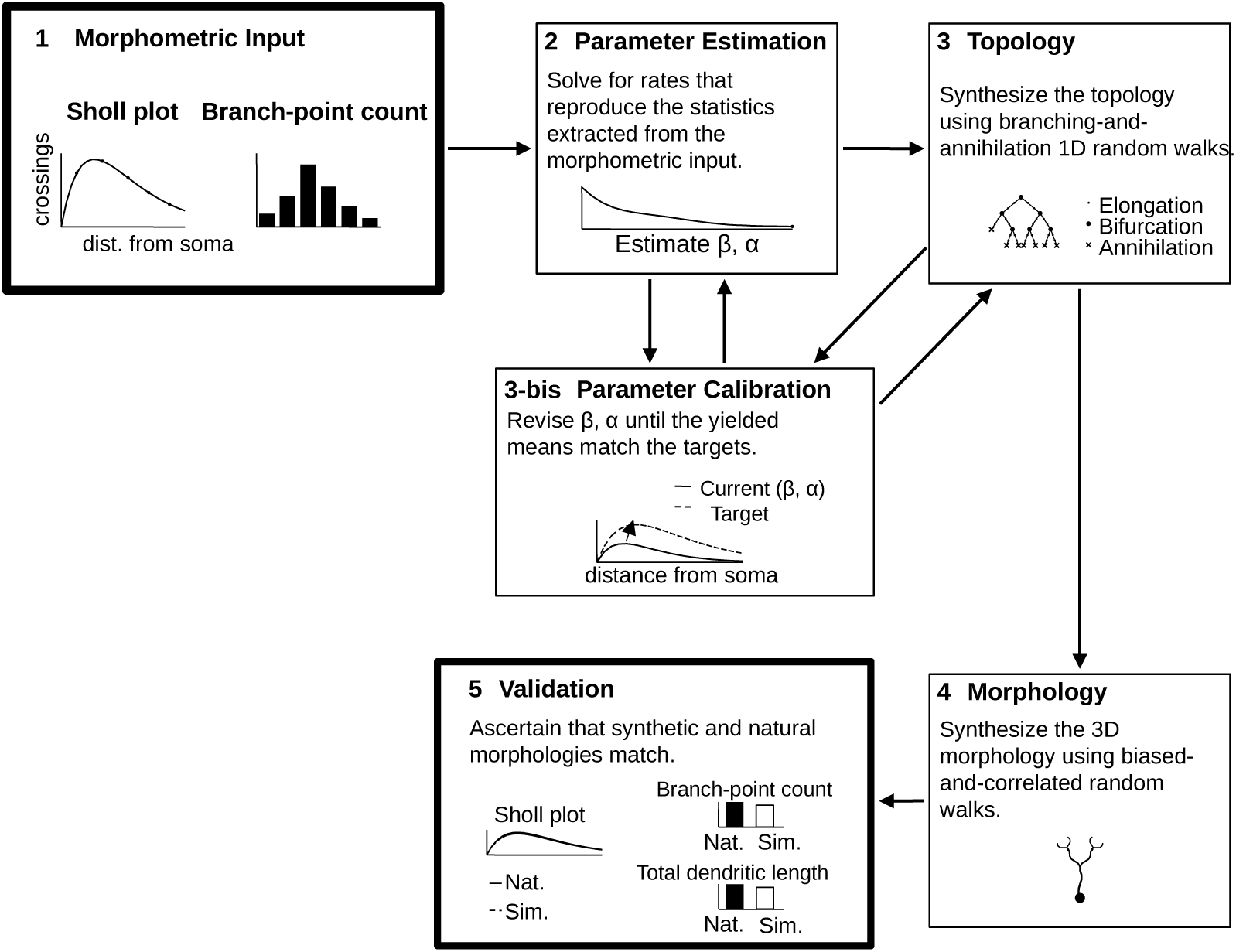
Overview of the NeuWalk synthesis pipeline. (1) Morphometric input: a Sholl plot and a branch-point count distribution extracted from experimental reconstructions. (2) Parameter estimation: branching (β) and annihilation (α) rates are solved for so as to reproduce the input statistics. (3) Topology: the calibrated rates drive mono-dimensional branching-and-annihilating random walks that synthesize the sequence of elongation, bifurcation, and annihilation events. (3-bis) Parameter calibration: since the variance-reduction procedure truncates the distributions and thereby shifts their means, the rates are revised iteratively, exchanging with parameter estimation (2) and topology synthesis (3), until the yielded mean Sholl intersections and branch-point counts match the experimental targets. (4) Morphology: the topology guides biased-and-correlated threedimensional random walks that synthesize the morphology. (5) Validation: synthetic morphologies are compared against their natural counterparts on Sholl profile, branchpoint count, and total dendritic length.

Bifurcation and annihilation rates are estimated from Sholl intersections, with bifurcation counts as an optional constraint, using closed-form equations introduced in our previous publication [31]. However, the original version of this method yielded topologies with excessive variance in Sholl intersections and bifurcation counts [31]. To solve this issue, we equipped the monodimensional random walk with a variance-reduction procedure that keeps both measures within k experimental standard deviations of the mean (see Section 3.3). Here, we set k to 3 to cover 99.7% of the experimental distribution, under the assumption of normality. However, this procedure truncates their distributions, thereby distorting mean values. To correct this issue, we developed a calibration procedure that revises the rates and realigns mean values for Sholl intersections and bifurcation counts (stage 3-bis; see Section 3.4).

Topologies prescribe occurrences of elongation, bifurcation, and annihilation in a three-dimensional random walk that synthesizes morphologies (stage 4; see Section 3.5), where user-specified combinations of biases sculpt patterns characteristic of natural morphologies (Figure 2). Finally, synthetic morphologies are validated by comparing Sholl profiles, bifurcation counts, and total dendritic length against estimates from natural counterparts (stage 5; see Section 3.7).

**Figure 2:**
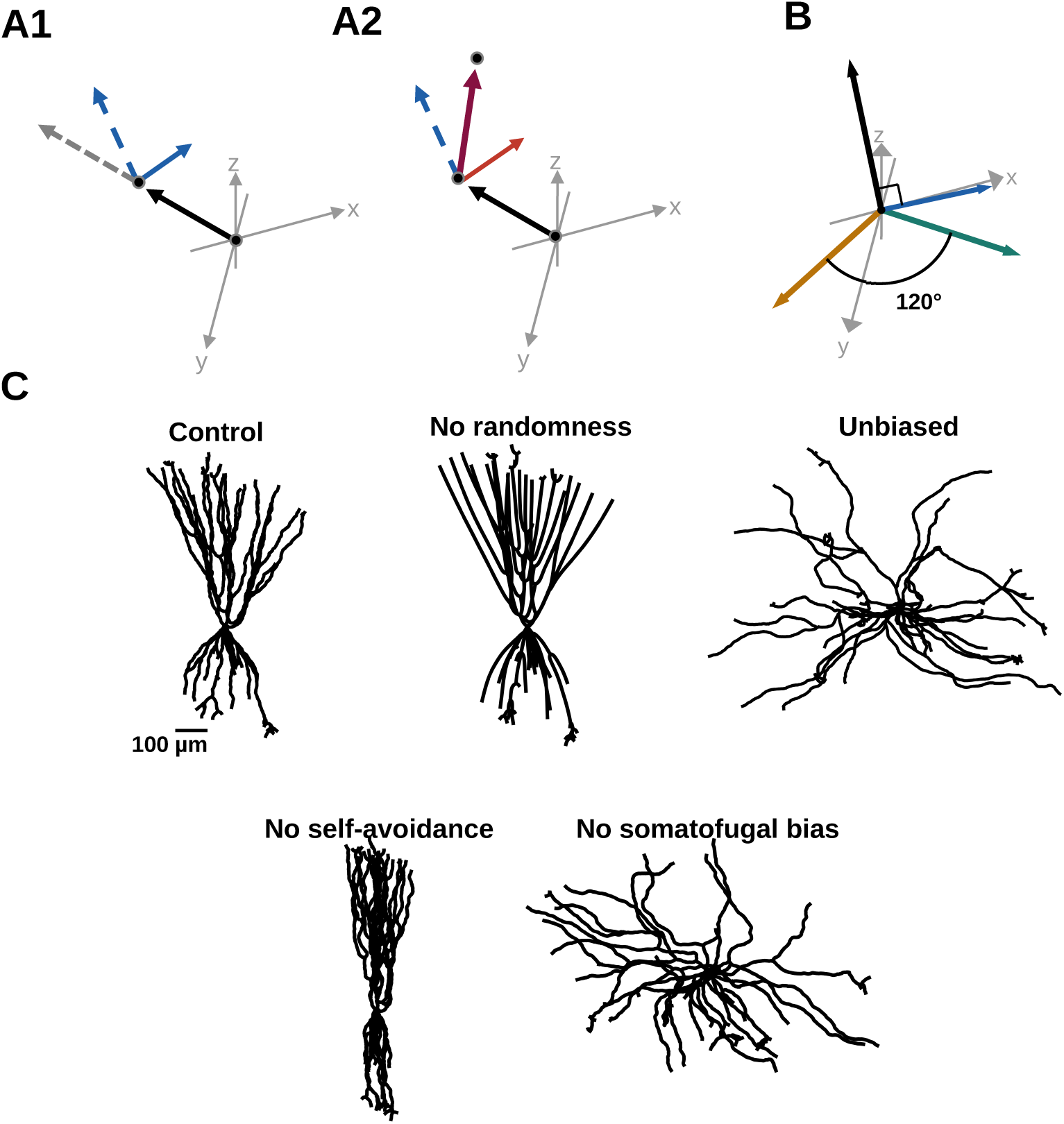
Mechanisms determining trajectory direction, and their contribution to synthetic morphology. (A) Direction update at each elongation step: the bias b⃗ is combined with the previous direction v⃗*_i_* via the sequential combination operator ⊗ (A1), and the random component r⃗ is subsequently combined with this result to give the new direction v⃗*_i_*_+1_ = v⃗*_i_*⊗ b⃗ ⊗ r⃗ (A2). (B) Bifurcation direction determination: the parent direction v⃗*_i_* and the aggregate repulsion R⃗ exerted by neighboring dendrites jointly determine a torsion about v⃗*_i_* that orients the bifurcation plane, within which the two offspring directions v⃗^(left)^_i+1_ and v⃗^(right)^_i+1_ are placed 120*^◦^* from v⃗*_i_* and from each other. (C) A synthetic pyramidal neuron (PYR) from the anterior piriform cortex under different model configurations, with all other parameters held fixed.

### 3.1. Curation of experimental morphological reconstructions

We extracted Sholl profiles, bifurcation counts, and total dendritic length from experimental reconstructions of middle tufted cells (mTCs; n = 5) and mitral cells (MCs; n = 9) from mouse main olfactory bulb [36]; semilunar cells (SLs; n = 3) and pyramidal neurons (PYRs; n = 12) from rat piriform cortex [15]; and neocortical pyramidal neurons of the thick-tufted subtype (NCPYRs; n = 10) from layer 5 of rat neocortex [2, 37]. Sholl profiles were computed with a bin size of 50 µm. All morphologies are publicly available on NeuroMorpho.Org [38].

Morphologies were curated prior to analysis. Specifically, we addressed three recurring issues: deformed or amputated dendritic trees, misplaced dendrites, and mislabeled dendrite types (for details, see Supplementary Materials). Axons were not a target of our study, as they were either absent or truncated in all reconstructions, thus making it impossible to obtain reliable estimates of morphometric measures. For mTCs and MCs, the apical/tuft dendritic complex was excluded from our analysis, as it appeared bent — lacking the natural orientation from the soma toward the glomerulus — missing, or misplaced. Yet synthesizing basal dendrites allowed us to capture the vast majority of dendrites in these cells, while fictional replacements of apical/tuft dendrites were synthesized for illustrative purposes only (see Figures 3 and 4). For SLs, basal dendrites were excluded from our analysis, as they were missing in one SL, leaving only two morphologies for the estimation of morphometric measures. However, apical dendrites account for the vast majority of dendritic length in these cells. Finally, the original reconstructions of NCPYRs did not distinguish between the apical trunk and oblique dendrites, so the latter were identified through visual inspection and labeled accordingly.

**Figure 3:**
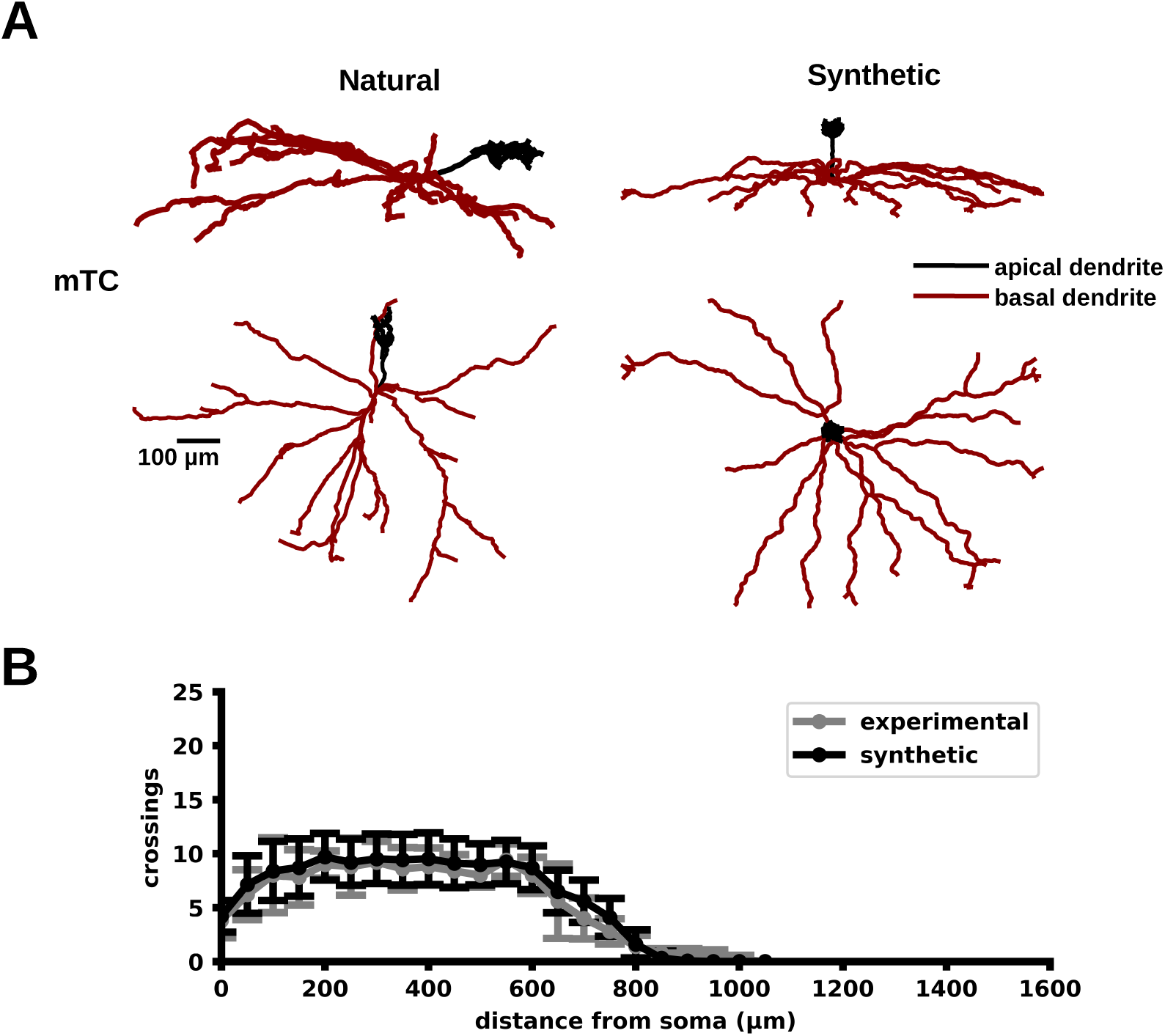
Validation for middle tufted cells from the olfactory bulb. (A) Sagittal (i.e., y-z axes; Top) and transverse (i.e., x-y axes; Bottom) views of natural (left) and synthetic (right) morphologies of middle tufted cells (mTCs). These cells have apical (black) and basal (red) dendrites. The apical dendrites of synthetic cells are illustrative only (for details, see Section 3.1). (B) Sholl plots of their basal dendrites across the sampled population (mean ± SD).

**Figure 4:**
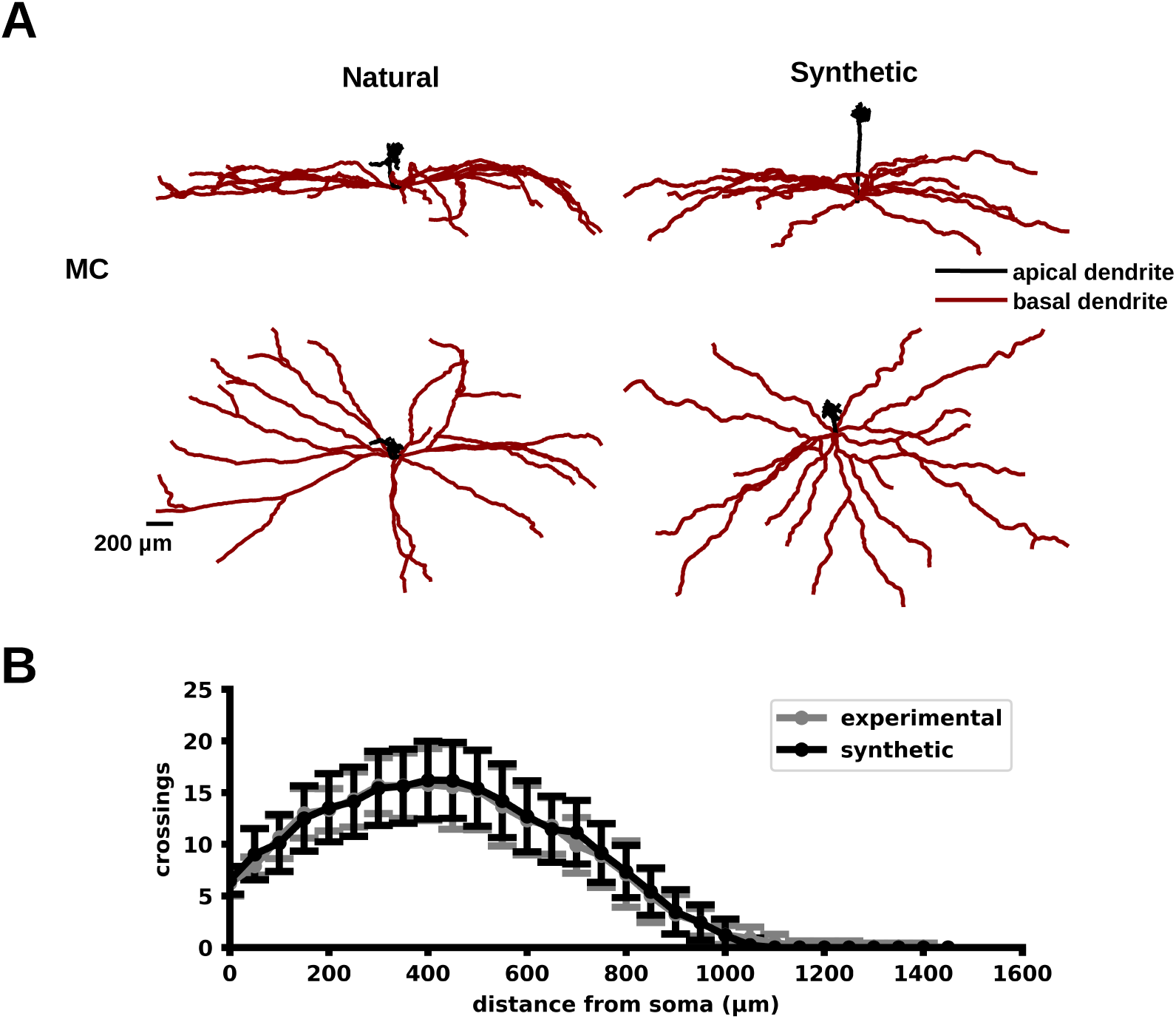
Validation for mitral cells from the olfactory bulb. Same as in Figure 3, for mitral cells (MCs).

### 3.2. Topology synthesis

Topology refers to a descriptor that indicates the sequence of elongation, bifurcation, and annihilation events that occur during the synthesis.

Our preliminary analysis of experimental reconstructions showed that the vast majority of segments were oriented centrifugally with respect to the soma (≫ 99%) [31], a property we called *somatofugality*. Motivated by these findings, we synthesize topologies using branching-and-annihilating mono-dimensional random walks, under the assumption that they drift away from the soma.

Topology synthesis begins by determining the number of primary dendrites (i.e., dendrites originating from the soma). Subsequently, at each step, a dendritic tip can undergo one of three mutually exclusive events: 1) *elongation*: the tip advances a distance equal to the step size Λ from the soma; 2) *bifurcation*: the tip bifurcates into two new tips, each advancing a distance Λ from the soma; and 3) *annihilation*: the tip terminates. Once a tip has bifurcated or annihilated, it cannot evolve further. The three events occur with probabilities p*_e_* (= 1 − p*_b_* − p*_a_*), p*_b_* (= βΛ), and p*_a_* (= αΛ), respectively, where p*_b_* and p*_a_* scale with the step size Λ according to the rates β and α, respectively.

Arrays of bifurcation (β*_i_*)*_i_*_=0*,···, n*_ and annihilation (α*_i_*)*_i_*_=0*,···, n*_ rates are directly estimated from Sholl intersections and bifurcation counts, the latter as an optional constraint, using closed-form equations introduced in our previous publication [31]. This yields a non-stationary branching-and-annihilating random walk, with rates estimated for each spatial interval spanning a Sholl bin. This formulation ensures that synthetic morphologies reproduce Sholl intersections and bifurcation counts in a way invariant with respect to the specific trajectories or spatial dimension of the random walk, as long as somatofugality is respected.

### 3.3. Variance-reduction procedure

To control the variance of Sholl intersections and bifurcation counts, we developed a progressive, bin-wise synthesis procedure with backtracking (see Algorithm 1), which confines the ranges of both measures in synthetic topologies to physiological boundaries. Specifically, branching-and-annihilating sequences are synthesized one Sholl bin at a time. If the number of Sholl intersections for the current bin falls within κ (= 3) experimental standard deviations of the mean, where κ is a tunable parameter, then it is accepted and recorded as a checkpoint. If a bin fails this test, the synthesis backtracks to the most recent checkpoint and the bin is re-synthesized. In addition, if repeated attempts at a given checkpoint fail, the procedure progressively expands its rollback window to include earlier, previously accepted bins, which are discarded and re-synthesized together with the current one. This allows the procedure to recover from local branching configurations that make the target number of intersections for a given bin statistically difficult to reach, without discarding the entire profile and restarting synthesis from the soma. Finally, the procedure verifies that the bifurcation count falls within κ experimental standard deviations of the mean. If the test fails, the entire topology is re-synthesized.

This procedure truncates the distributions of Sholl intersections and bifurcation counts in synthetic topologies, thereby shifting their mean values. To correct this issue, we developed the calibration procedure, described in the next section.

### 3.4. Parameter calibration procedure

Topology synthesis is a non-stationary Galton–Watson process that tends to yield right-skewed distributions of Sholl intersections and bifurcation counts. Thus, the variance-reduction procedure truncates these distributions, thereby shifting their mean values. To ensure that the mean values are reproduced by the synthesis, we developed a calibration procedure that iterates topology synthesis (see Algorithm 2). Here we distinguish between three mean values:

- the *experimental mean*, estimated directly from Sholl intersections and bifurcation counts;
- the *synthetic mean*, yielded by the topology synthesis;
- the *parameter mean*, provided as input to the topology synthesizer.

The procedure iteratively shifts the parameter means for the Sholl profile and the bifurcation count until the difference between synthetic and experimental means falls below the tolerances τ*_S_* (= 1) and τ*_B_* (= 1), fixed a priori for Sholl intersections and bifurcation counts, respectively. Specifically, the procedure calibrates one Sholl bin at a time, proceeding from proximal to distal bins. For each bin, the topology is re-synthesized, and the synthetic mean of intersections is tested against the corresponding experimental mean. If the discrepancy falls below τ*_S_*, the procedure moves on to the next bin; otherwise, the parameter mean is shifted by the observed difference and the topology is re-synthesized and tested. If convergence does not occur within the iteration budget I (= 20), failure is declared for that bin and the procedure moves to the next one. The bifurcation count is then calibrated the same way, using topologies synthesized to their full extent rather than up to a single bin, with failure declared under the same condition and iteration budget.

Note that calibration works in two steps: parameter means for Sholl intersections are calibrated first, followed by those for bifurcation counts. This order was chosen because preliminary tests indicated that Sholl profiles are robust across a wide range of bifurcation counts. After calibration, the corrected parameter means are used to synthesize the topologies that guide morphology synthesis, described in the next Section.

### 3.5. Morphology synthesis

We synthesize morphologies using biased-and-correlated three-dimensional random walks, following the sequences of elongation, bifurcation, and annihilation events prescribed by topological descriptors.

This process begins at the soma by setting the initial orientations of the primary dendrites. At any given iteration, each tip extends along a random trajectory guided by combinations of biases, each contributing a specific directional pattern, which together sculpt the distinctive features observed in natural morphologies. By leveraging their intuition, users infer a combination of biases from images of neuron morphologies or qualitative descriptions thereof. In this respect, tips mimic the behavior of growth cones, the motile structures at the end of developing neurites, where biases act as phenomenological models of the guidance cues that direct neurite extension. The random walk is correlated because each new direction is derived from the preceding one, rather than being drawn independently at every step. This confers directional persistence, yielding trajectories that turn gradually, as observed in dendrites, rather than erratically. Moreover, the tortuosity of the resulting dendrites arises from the interplay between the intensity of the random component of the walk and that of the biases.

Directions of dendritic tips for elongation and bifurcation are governed by different bias functions. In both cases, a step displaces the tip by a length λ, corrected at each iteration so that the tip advances by a fixed distance Λ, independently of the exact direction, thus reproducing exactly the somatofugal drift assumed during topological synthesis.

#### 3.5.1. Elongation

At each elongation step, the position of the new tip p*_i_*_+1_ is computed as

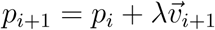

where p*_i_* is the last tip position, v⃗*_i_*_+1_ is a unit vector indicating the extending direction of the tip, and λ is the step length. In particular, v⃗*_i_*_+1_ is obtained by combining the last direction v⃗*_i_* with one or more vectors b⃗*_j_* representing different biases, along with a vector r⃗, representing a random component, using the sequential combination ⊗, such that

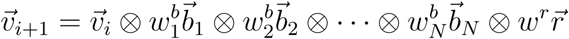

where *w^b^_j_* and w*^r^* are non-negative scalar values that set the relative contribution of each vector to the direction v⃗*_i_*_+1_. In particular, the sequential combination ⊗ returns a unit vector by normalizing the sum of its two arguments (Figure 2A1 and A2). This operator is commutative but not associative, so that the application order of biases matters. Moreover, bias functions can be derived by using linear combinations such that

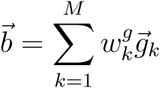

where b⃗ is the derived bias and g⃗*_k_*are biases calculated by specific functions.

The user is therefore expected to select biases *(b⃗_j_)_j=1,…,N_* together with their weights and the weight of the random component r⃗.

Each elongation bias can be configured such that its intensity scales with the distance from its source (e.g., root, dendrite, or surface). Such a dependence is modeled by a Hill-type function:

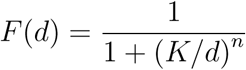

where d is a bias-specific distance measured from the corresponding source and evaluated at the current tip position, K is the half-maximal distance and n the Hill coefficient. In particular, a positive n yields a bias that strengthens with distance, whereas a negative n yields one that weakens with it.

Moreover, each elongation bias contributes a distinct morphological pattern, as shown by ablating individual components from a synthetic pyramidal neuron of the anterior piriform cortex (Figure 2C, Control). For example, disabling self-avoidance alone results in fasciculation (No self-avoidance); disabling the somatofugal bias alone removes the somatofugal drift that orients dendrites away from the soma (No somatofugal bias). Together, these ablations indicate that specific bias combinations control complementary aspects of morphology, each being necessary to reproduce realistic dendritic organization. Elongation biases implemented in the package, along with their aliases, are described in Table 1.

**Table 1:** Elongation bias functions registered in neuwalk.synthesis.morphology.biases.elongation, identified by the alias passed to biases.get elongation(alias, …).

| Alias | Description |
| --- | --- |
| `plane_boundary` | Pulls growth toward (or, with **resistance**, away from once already aligned) a given plane. |
| `attraction` | Pulls growth toward a fixed target point, with optional distance-dependent strength. |
| `truncated_cone_boundary` | Pulls growth toward the central axis of a truncated cone, whose cross-sectional radius can taper along its length. |
| `ellipsoid_boundary` | Pulls growth toward (inward) or away from (outward) the surface of an ellipsoid, confining dendrites within or outside a curved boundary. |
| `sibling_repulsion` | Pushes a section away from its own sibling sections (sharing the same parent). |
| `parent_repulsion` | Pushes a section away from its own parent section specifically. |
| `all_sections_repulsion` | Pushes a section away from every other section in the whole tree (excluding soma). |
| `nonrelated_repulsion` | Pushes a section away from other sections in the same root's tree, excluding its own parent and siblings. |
| `root_repulsion` | Pushes a section radially outward from its own root's first point (soma-attachment point). |

#### 3.5.2. Bifurcation

A bifurcation bias computes the initial directions of the two offspring branches from the last direction of the parent (Figure 2B). Given the last direction v⃗^(parent)^_last_ of the parent at the bifurcation, a torsion R⃗ — a rotation about it — determines the lateral component for the directions of the off-spring as *l^ = (v⃗^(parent)^_last_ × R⃗)/‖v⃗^(parent)^_last_ × R⃗‖*. Thus, the directions of the offspring follow as

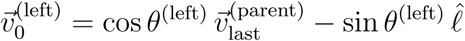

and

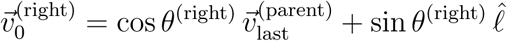

where θ^(left)^ and θ^(right)^ are the bifurcation angles of the two offspring. This method makes the parent and offspring directions coplanar [39], with the torsion perpendicular to the bifurcation plane. When both angles are set to 60*^◦^*, each pair of directions forms 120*^◦^* angles [40].

As for elongation, the user is expected to select a bifurcation bias by leveraging intuition. Unlike elongation biases, bifurcation biases cannot be combined through sequential or linear combinations. Bifurcation biases implemented in the package, along with their aliases, are described in Table 2.

**Table 2:** Bifurcation bias functions registered in neuwalk.synthesis.morphology.biases.bifurcation, identified by the alias passed to biases.get bifurcation(alias, …).

| Alias | Description |
| --- | --- |
| `radial_torsion` | Splits the two offspring symmetrically at a fixed angle from the parent's own direction, on a bifurcation plane whose normal is the aggregate repulsion from all other sections, so that the split twists away from crowded neighbours. |
| `internal_branch` | One offspring continues straight along the parent's own last direction; the other branches off at a fixed angle with a random azimuthal orientation. |
| `cross_torsion` | Same symmetric-split mechanism as `radial_torsion`, but the normal of the bifurcation plane is the surface normal of an ellipsoid (or unit sphere) at the bifurcation point, instead of section repulsion. |

### 3.6. Subclass-specific rate estimation

Based on differing branching patterns and relative orientation, NeuWalk divides apical dendrites into trunk and obliques, using distinct configurations for their synthesis.

Specifically, it estimates the spatial distribution of the bifurcation points at which oblique trees branch off the apical trunk, together with the number of oblique trees per neuron. It then detaches the oblique trees from the apical trunk, translates them to the origin, and analyzes them as if they were a whole dendritic tree with a common root. The bifurcation points of obliques are no longer counted for the apical trunk. Subsequently, it synthesizes the topologies of the apical trunk and the obliques separately, and then attaches the oblique topologies to that of the apical trunk, drawing their number and their grafting sites from the aforementioned distributions.

Morphological synthesis proceeds in a different order. Specifically, oblique morphologies are synthesized from the trunk outward: the apical trunk is synthesized first, and each oblique subtree then extends from its grafting site, using different configurations of biases for the two types of dendrites.

Among the neuron classes analyzed here, this strategy was applied to NCPYRs, the only class bearing oblique dendrites.

### 3.7. Morphology validation

We validated synthetic morphologies by comparing Sholl intersections, bifurcation counts, and total dendritic lengths against their natural counterparts, using a two-tailed bootstrap hypothesis test introduced in our previous publication [31]. Sholl intersections were compared on a bin-by-bin basis (bin width: 50 µm; see Supplementary Tables S1–S5 and S8). This non-parametric approach allowed running statistical tests on a limited pool of experimental reconstructions. Each synthetic population comprised 200 morphologies. Synthetic and natural morphologies were resampled 10,000 times to construct empirical null distributions of mean and variance differences between groups and thus compute p-values. Statistical significance was assessed at α = 0.01. To compare the number of Sholl intersections, a Bonferroni correction was applied. A minimum-effect-size threshold was applied alongside the bootstrap significance test, since statistical detectability and practical importance are distinct [41, 42]: a difference was flagged as meaningful only if it was both significant and exceeded a minimum relative mean difference (10%) or variance ratio (1.5), as applicable. When a bin’s variance was exactly zero for one or both groups, the variance-ratio test was not run, as the comparison is undefined in this case. To remove the dependence of the variance-ratio test’s p-value on the order of the two groups, the reject and-redraw handling of degenerate (zero-variance) bootstrap resamples was replaced by adding a symmetric floor to every resampled variance. Comparisons were only performed when each group had more than two neurons. The analysis was repeated for the basal and apical dendritic trees of each neuron class.

### 3.8. Software performance testing

NeuWalk was tested on a workstation equipped with an Intel Core i9-11900H processor operating at 2.5–4.9 GHz and 32 GB of DDR4-3200 MHz RAM. To assess the computational efficiency and practical usability, we estimated the time required for each synthesis, which ranged from 10 to 40 seconds per morphology.

## 4. Results

### 4.1. Olfactory bulb: mitral and tufted cell synthesis

We tested NeuWalk by synthesizing MCs and mTCs (Figures 3A and 4A). These neurons comprise apical/tuft (black) and basal dendrites (red). Our analysis focused on basal dendrites, which account for ∼87% of total dendritic length; apical/tuft dendrites were synthesized for illustrative purposes only (for details, see Section 3.1).

To synthesize basal dendrites, we began by approximating the external plexiform layer (EPL) boundaries as two ellipsoidal surfaces: an outer one (axes: 2000 × 1250 × 1250 µm), corresponding to the glomerular layer, and an inner one (150 µm beneath it) [43]. Within this volume, basal dendrites of both cell types extend at different depths (mTCs: 75–150 µm; MCs: 0– 75 µm; calculated from the inferior surface). The surfaces acted as sources for two ellipsoid boundary biases, both with distance dependence (n = −1): the inferior and superior boundaries exerted outward (K = 1 µm for both cell types) and inward (K = 50 µm and K = 75 µm for mTCs and MCs, respectively) repulsions. Together, these confined basal dendrites to the sublamina of each cell type, while conferring the characteristic curvature observed in anatomical studies (Figures 3A and 4A, top) [43, 36]. Also, the root repulsion bias realized somatofugality, conferring the radial patterns (Figures 3A and 4A, bottom). Self-avoidance resulted from a linear combination of sibling repulsion and nonrelated repulsion biases, both distance-dependent (K = 20 µm and K = 10 µm, respectively; n = −2 for both). As bifurcation bias, we used cross torsion, which kept the bifurcation plane parallel to the ellipsoidal boundaries of the EPL.

For both mTCs and MCs, mean and variance of Sholl intersections (Figures 3B and 4B; per-bin comparisons in Supplementary Tables S1 and S2), bifurcation count (Table 3), and total dendritic length (Table 4) were statistically indistinguishable between synthetic and natural morphologies.

**Table 3:** Bifurcation count, natural (Nat) vs. synthetic (Synth) populations, reported as mean ± SD. n denotes the number of natural reconstructions for that cell type. p_mean_ and p_var_ are two-tailed bootstrap p-values (see Section 3.7; 10,000 resamples, α = 0.01) for the difference in mean and in variance, respectively; asterisks denote differences that were both significant and above the minimum effect size (see Section 3.7). Arbor type is reported separately where dendritic subtypes are modeled independently (see Section 3.6). Apical/tuft dendrites of mTCs and MCs were not synthesized in a data-driven manner, so only their natural values are reported (see Section 3.1). *^†^*Basal dendrites of SLs were excluded from synthesis and analysis, as they were missing in one reconstruction, leaving only two morphologies for parameter estimation; their natural values are reported for reference (see Section 3.1).

| Cell type | Arbor | $n$ | Nat | Synth | $p_{\text{mean}}$ | $p_{\text{var}}$ |
| --- | --- | --- | --- | --- | --- | --- |
| mTC | Basal | 5 | $16.0 \pm 3.1$ | $16.3 \pm 3.4$ | 0.812 | 0.877 |
| mTC | Apical/tuft | 5 | $43.2 \pm 9.6$ | — | — | — |
| MC | Basal | 9 | $17.4 \pm 6.0$ | $16.8 \pm 6.6$ | 0.756 | 0.878 |
| MC | Apical/tuft | 9 | $27.6 \pm 17.1$ | — | — | — |
| SL | Basal<sup>†</sup> | 2 | $16.0 \pm 8.7$ | — | — | — |
| SL | Apical | 3 | $22.0 \pm 4.3$ | $22.2 \pm 6.7$ | 0.894 | 0.337 |
| PYR | Basal | 12 | $16.0 \pm 8.7$ | $15.7 \pm 10.9$ | 0.899 | 0.381 |
| PYR | Apical | 12 | $18.8 \pm 4.3$ | $18.3 \pm 7.2$ | 0.686 | 0.181 |
| NCPYR | Basal | 10 | $29.2 \pm 12.3$ | $29.5 \pm 15.0$ | 0.947 | 0.480 |
| NCPYR | Apical | 10 | $53.3 \pm 9.5$ | $54.0 \pm 15.5$ | 0.822 | 0.118 |

**Table 4:**
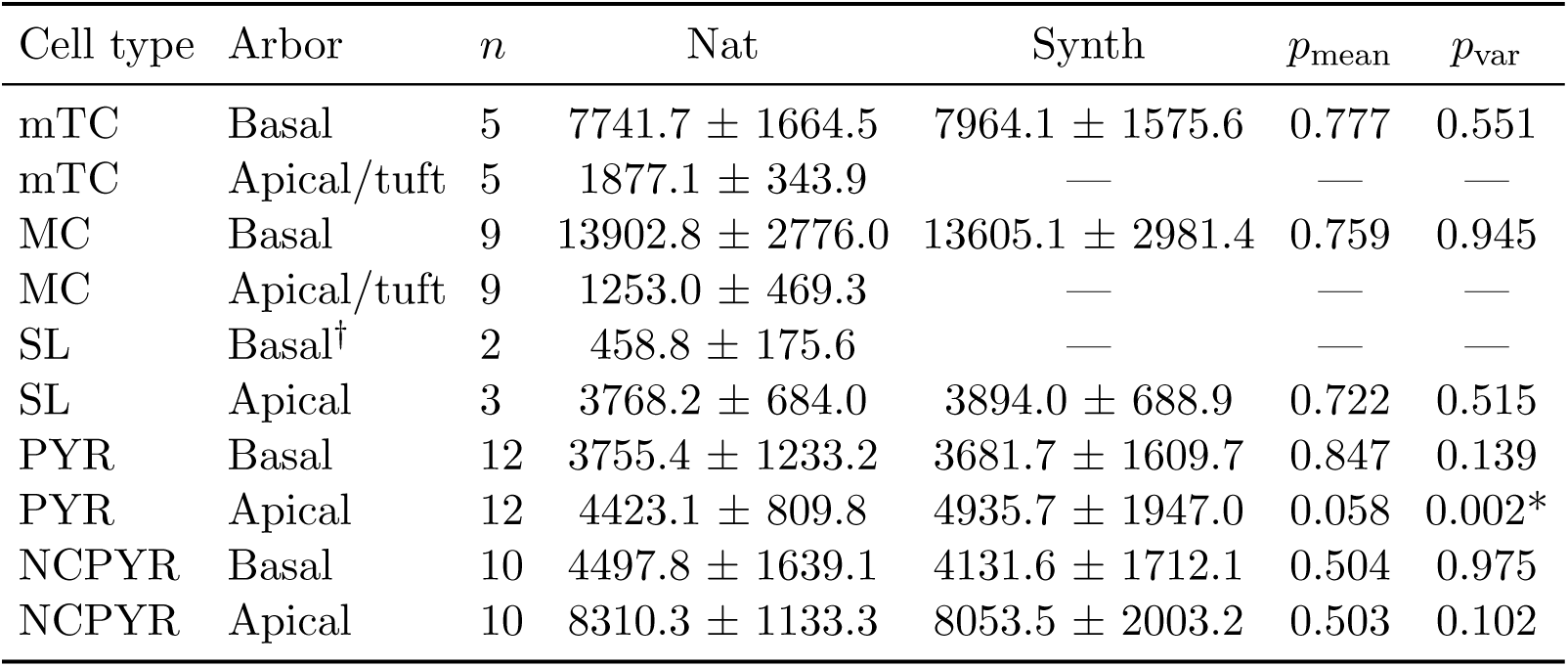
Total dendritic length (µm), natural (Nat) vs. synthetic (Synth) populations, reported as mean ± SD. n denotes the number of natural reconstructions for that cell type. p_mean_ and p_var_ are two-tailed bootstrap p-values (see Section 3.7; 10,000 resamples, α = 0.01) for the difference in mean and in variance, respectively; asterisks denote differences that were both significant and above the minimum effect size (see Section 3.7). Arbor type is reported separately where dendritic subtypes are modeled independently (see Section 3.6). Apical/tuft dendrites of mTCs and MCs were not synthesized in a data-driven manner, so only their natural values are reported (see Section 3.1). *^†^*Basal dendrites of SLs were excluded from synthesis and analysis, as they were missing in one reconstruction, leaving only two morphologies for parameter estimation; their natural values are reported for reference (see Section 3.1).

| Cell type | Arbor | $n$ | Nat | Synth | $p_{\text{mean}}$ | $p_{\text{var}}$ |
| --- | --- | --- | --- | --- | --- | --- |
| mTC | Basal | 5 | $7741.7 \pm 1664.5$ | $7964.1 \pm 1575.6$ | 0.777 | 0.551 |
| mTC | Apical/tuft | 5 | $1877.1 \pm 343.9$ | — | — | — |
| MC | Basal | 9 | $13902.8 \pm 2776.0$ | $13605.1 \pm 2981.4$ | 0.759 | 0.945 |
| MC | Apical/tuft | 9 | $1253.0 \pm 469.3$ | — | — | — |
| SL | Basal<sup>†</sup> | 2 | $458.8 \pm 175.6$ | — | — | — |
| SL | Apical | 3 | $3768.2 \pm 684.0$ | $3894.0 \pm 688.9$ | 0.722 | 0.515 |
| PYR | Basal | 12 | $3755.4 \pm 1233.2$ | $3681.7 \pm 1609.7$ | 0.847 | 0.139 |
| PYR | Apical | 12 | $4423.1 \pm 809.8$ | $4935.7 \pm 1947.0$ | 0.058 | 0.002* |
| NCPYR | Basal | 10 | $4497.8 \pm 1639.1$ | $4131.6 \pm 1712.1$ | 0.504 | 0.975 |
| NCPYR | Apical | 10 | $8310.3 \pm 1133.3$ | $8053.5 \pm 2003.2$ | 0.503 | 0.102 |

### 4.2. Piriform cortex: semilunar and pyramidal cell synthesis

We further tested our method on SLs and PYRs (Figures 5A and 6A). These neurons comprise apical (black) and basal dendrites (red; not shown for SLs). Our analysis focused on both dendrite types for PYRs, whereas it focused on apical dendrites only for SLs (for details, see Section 3.1).

**Figure 5:**
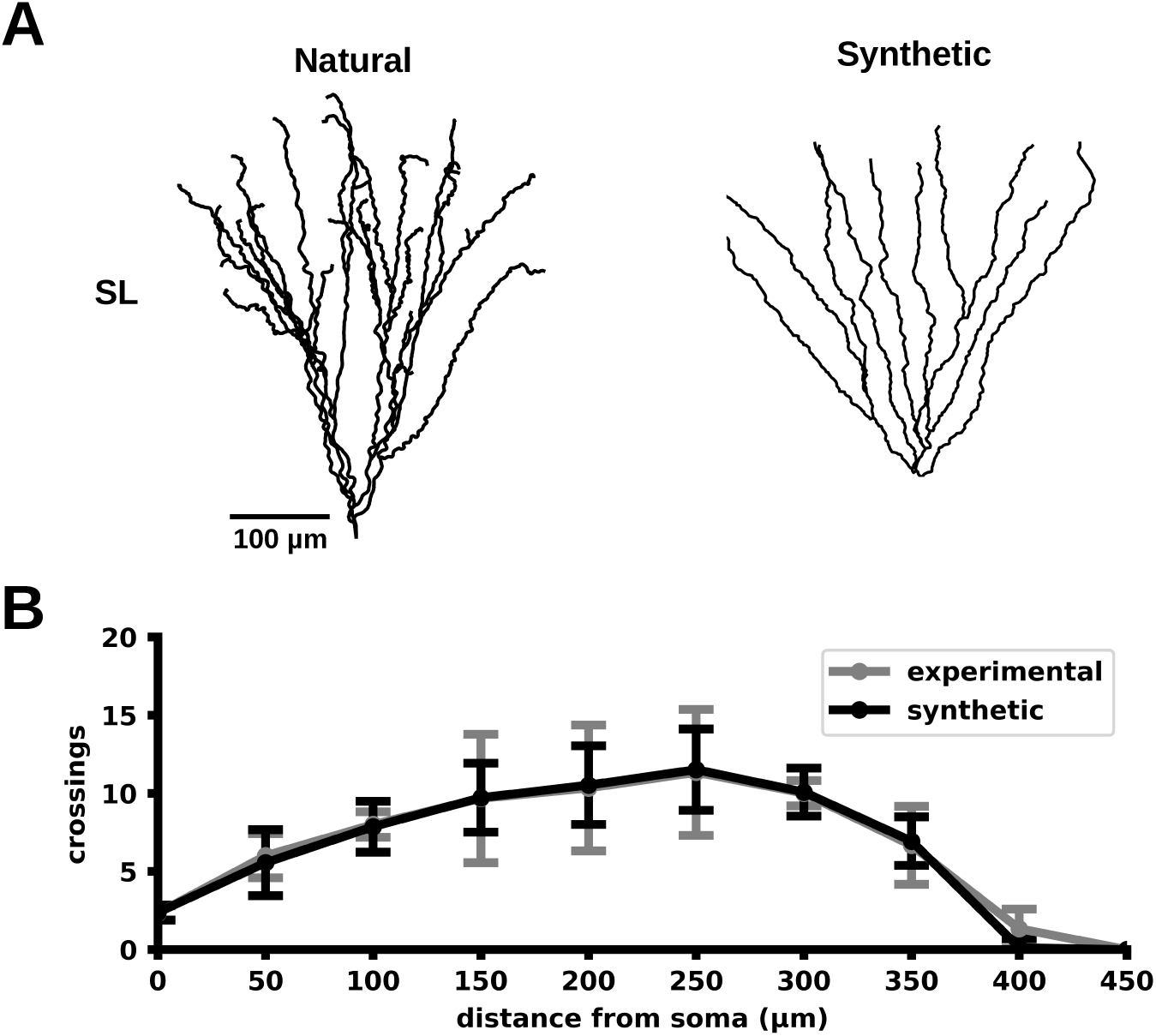
Validation for semilunar cells from the anterior piriform cortex. (A) Representative morphologies of natural (left) and synthetic (right) semilunar cells (SLs). Only apical dendrites (black) were analyzed (for details, see Section 3.1). (B) Sholl plot of their apical dendrites across the sampled population (mean ± SD).

**Figure 6:**
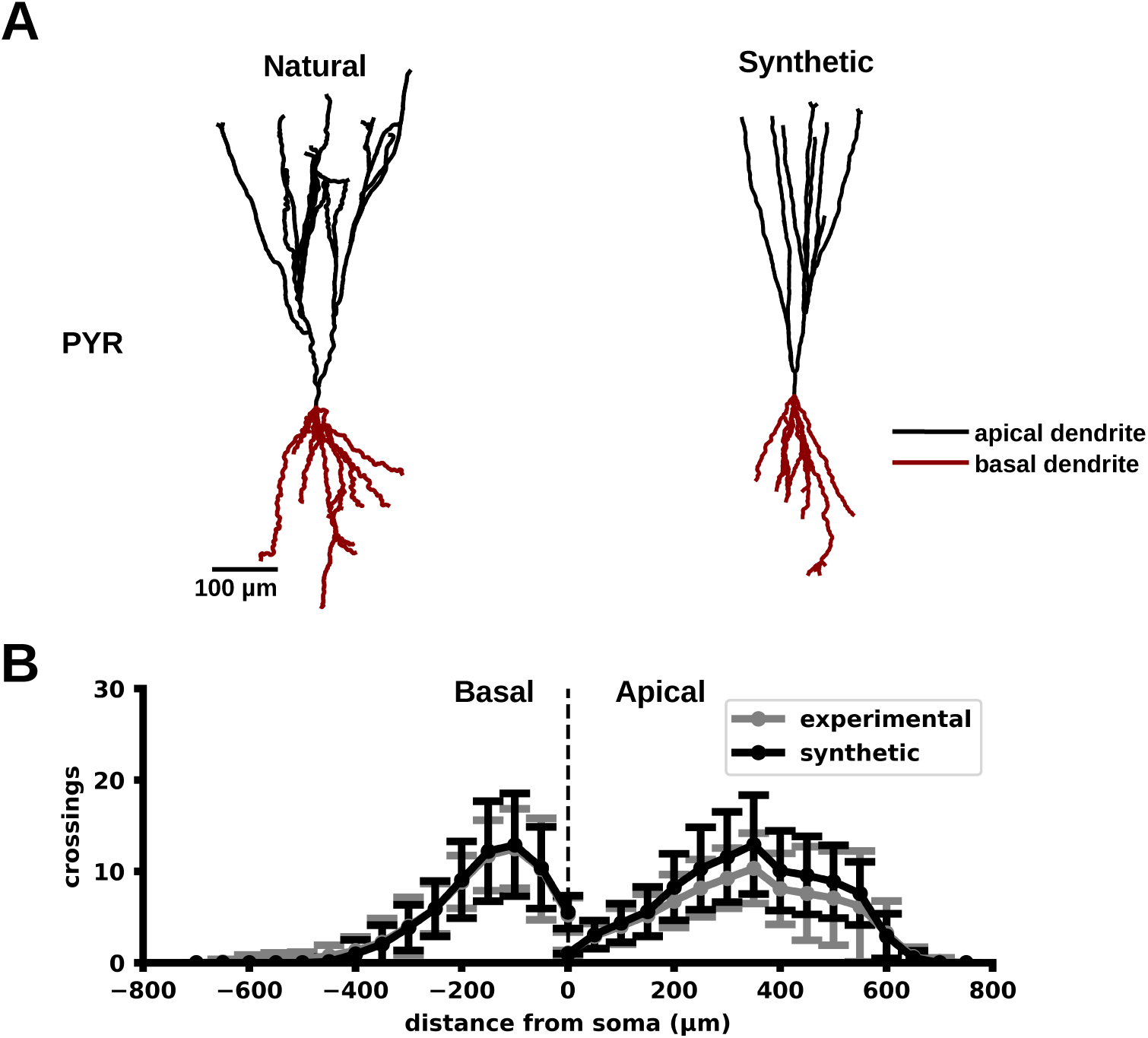
Validation for pyramidal neurons from the anterior piriform cortex. (A) Representative morphologies of natural (left) and synthetic (right) pyramidal neurons (PYRs). Unlike semilunar cells, both basal (red) and apical (black) dendrites were analyzed. (B) Sholl plots of their basal and apical dendrites across the sampled population (mean ± SD).

Both cell types share the same elongation and bifurcation biases. Two parallel plane boundary biases confined dendrites within a slab (of 25 µm and 50 µm for SLs and PYRs, respectively), repelling them in a distance-dependent fashion (K = 12.5 µm and K = 25 µm for SLs and PYRs, respectively; n = −2 for both). Another plane boundary bias repelled basal and apical dendrites downward and upward, respectively, creating an orientation along the somatodendritic axes as a form of somatofugality, in a distance-dependent fashion (K = 200 µm and K = 400 µm for apical dendrites of SLs and PYRs, respectively; K = 200 µm for basal dendrites of PYRs; n = −2 for all). Self-avoidance resulted from sibling repulsion and nonrelated repulsion biases in the same configurations as for mTCs and MCs. As bifurcation bias, we used radial torsion, with the torsion resulting from the combined repulsions of all dendrites.

For both SLs and PYRs, mean and variance of Sholl intersections (Figures 5B and 6B; per-bin comparisons in Supplementary Tables S3 and S4), bifurcation count (Table 3), and total dendritic length (Table 4) were statistically indistinguishable between synthetic and natural morphologies, except for the variance of total dendritic length in PYR apical dendrites.

### 4.3. Neocortex: pyramidal neuron synthesis

We finally tested our method on NCPYRs (Figure 7A). These neurons comprise an apical trunk (black), obliques (green), and basal dendrites (red). Similarly to SLs and PYRs, two parallel plane boundary biases confined dendrites within a slab (of 100 µm), exerting a distance-dependent repulsion (K = 50 µm and n = −2), while another plane boundary bias centered 1000 µm from the soma bent distal apical dendrites by exerting distance-dependent repulsion (K = 100 µm and n = −2). A third plane boundary bias repelled apical dendrites upward with distance-dependent intensity (K = 800 µm and n = −2), while the root repulsion bias repelled basal dendrites radially with distance-dependent intensity (K = 200 µm and n = −2), both realizing somatofugality. In addition, the same bias repelled each oblique subtree from its bifurcation site on the apical trunk with uniform intensity. Self-avoidance was defined as for SLs and PYRs. As bifurcation bias, the same radial torsion bias was used for apical trunk, basal, and oblique dendrites, while at the bifurcation sites where oblique dendrites grafted onto the apical trunk, the internal branch bias was used.

**Figure 7:**
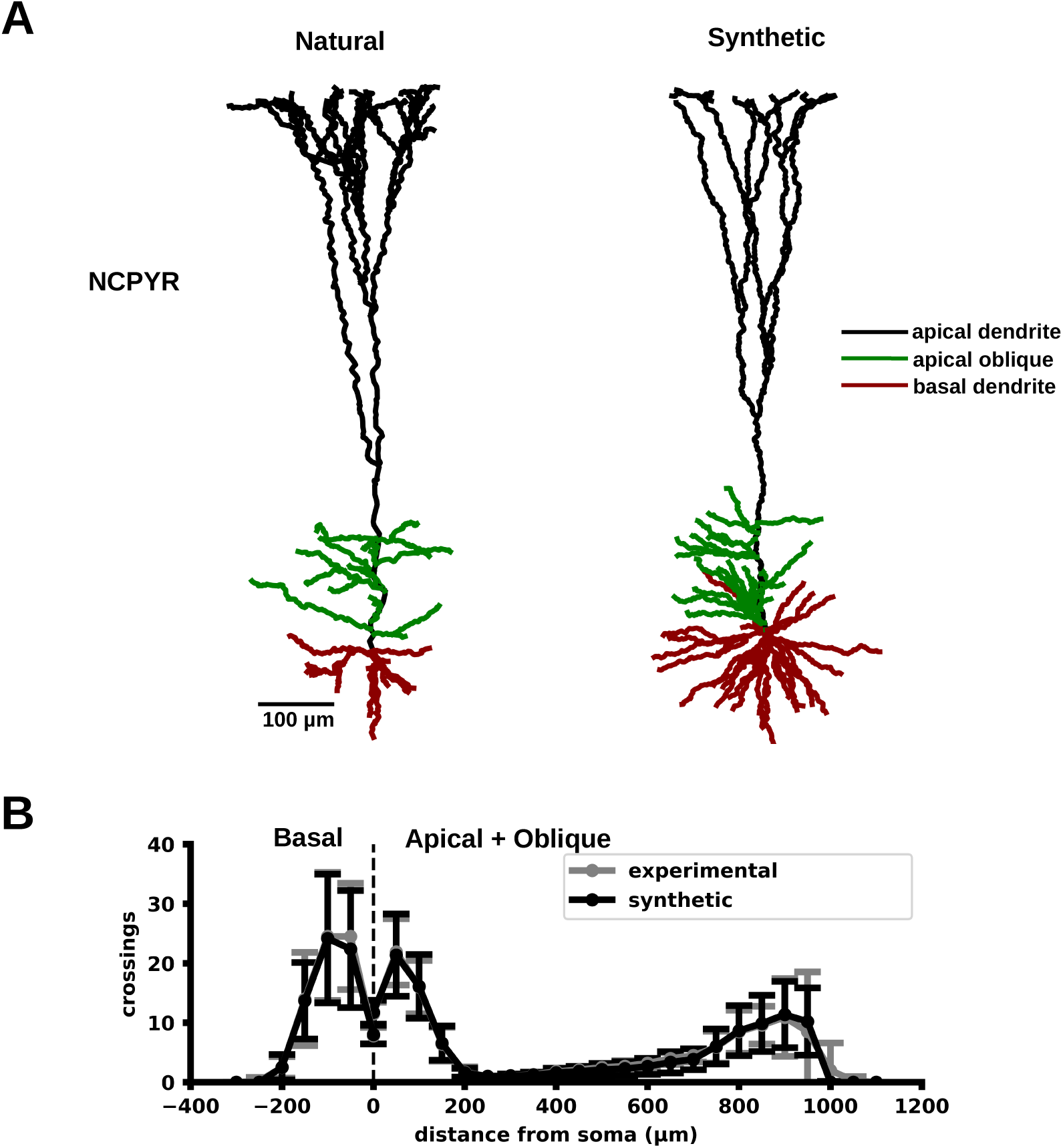
Validation for neocortical pyramidal neurons. (A) Representative morphologies of natural (left) and synthetic (right) neocortical pyramidal neurons (NCPYRs). These cells have basal dendrites (red), a main apical trunk (black), and obliques (green). (B) Sholl plots of their basal dendrites and of the combined apical and oblique arbor across the sampled population (mean ± SD).

For NCPYRs, mean and variance of Sholl intersections (Figure 7B; per-bin comparisons in Supplementary Table S5), bifurcation count (Table 3), and total dendritic length (Table 4) were statistically indistinguishable between synthetic and natural morphologies. Likewise, apical trunk and obliques showed no significant differences when tested separately (Tables S6–S8).

## 5. Discussion

We introduced NeuWalk, a Python library that synthesizes realistic neuron morphologies using branching-and-annihilating, biased-and-correlated random walks, validating it on five neuron classes from three brain regions. The validation of synthetic morphologies showed that the mean and variance of Sholl intersections, bifurcation counts, and total dendritic length were statistically indistinguishable from their natural counterparts, with a single exception discussed below.

Replication of Sholl intersections and bifurcation counts is a property of the method rather than an outcome of parameter search. Specifically, bifurcation and annihilation rates are estimated from these statistics, while variance-reduction and calibration procedures keep the resulting means and variances within physiological ranges, ensuring that targeted measures are replicated by construction. In contrast, total dendritic length is the only emergent property that depends on both the topology and the three-dimensional trajectories, therefore providing an independent check on the entire process.

The one exception concerns exactly this emergent measure. For apical dendrites of pyramidal neurons from the anterior piriform cortex, synthetic morphologies were about twice as variable in total dendritic length as their natural counterparts (SD = 1947 against 810 µm). Since the natural population comprises only twelve reconstructions, its variance is itself estimated with considerable uncertainty. On the other hand, the fact that variance was higher indicates that the method still spans the full range of variability, although a variance-reduction procedure specific to total dendritic length may become necessary if further tests confirm the same issue in other neuron types.

Compared with the approaches reviewed above, NeuWalk shifts the information requirement rather than removing it. All the reviewed methods require immediate access to morphological reconstructions to estimate parameters, whereas NeuWalk requires only Sholl intersections and, optionally, bifurcation counts, which are routinely reported in anatomical studies. Regarding the empiricism that other methods place in parameter fitting, in NeuWalk it is relocated to the selection of biases, which users can infer from published images or qualitative descriptions. Moreover, in NeuWalk structural errors cannot occur by construction. For example, since every branch bifurcates into exactly two offspring branches, and the number of primary dendrites coincides with the count of the first Sholl bin, it is impossible to obtain nodes with implausibly high branching degree as observed with MorphVAE.

Validation here is purely morphometric, without addressing whether synthetic morphologies support realistic electrotonic and integrative properties, which will be the goal of future work. This step will be pivotal for the reconstruction of large-scale models of the brain, requiring modeling ion channels while accounting for their kinetic properties and subcellular distributions, as done in previous modeling studies [3, 2]. This will allow us to explore the implications of the patterns emerging from the synthesis for neuronal function, and to ascertain whether synthetic single-neuron models constructed from our morphological synthesizer are biologically plausible, both individually and in their contribution to network dynamics. Finally, we plan to train artificial intelligence or machine learning models either to synthesize morphologies directly or to correct those already synthesized.

## 6. Conclusion

NeuWalk is a new Python library based on branching-and-annihilating, biased-and-correlated random walks, which enables the synthesis of realistic three-dimensional neuron morphologies. This library uses Sholl intersections and bifurcation counts as constraints, thereby ensuring their replication in synthetic morphologies, while the user empirically defines a set of biases that shape the characteristic patterns of each morphological class.

Morphology synthesizers are fundamental components in constructing large-scale biophysically-detailed models of the brain to obtain morphologically realistic populations. As Sholl intersections and bifurcation counts are commonly reported in the literature, NeuWalk can operate without requiring immediate access to experimental morphological reconstructions.

## CRediT authorship contribution statement

**Francesco Cavarretta:** Conceptualization, Methodology, Software, Validation, Formal analysis, Investigation, Data curation, Writing – original draft, Writing – review & editing, Visualization, Project administration.

## Declaration of competing interest

The author declares that he has no known competing financial interests or personal relationships that could have appeared to influence the work reported in this paper.

## Acknowledgements

The author dedicates his work to the memory of his father, whose example remains a lasting source of inspiration. He also thanks his wife for her constant support, along with Mimi and Juju for their steady companionship throughout the writing of this manuscript.

## Funding

This research did not receive any specific grant from funding agencies in the public, commercial, or not-for-profit sectors.

## Data availability

The source code of NeuWalk, together with the curated experimental morphologies and the synthetic morphologies used for validation, is publicly available on GitHub at https://github.com/fcavarretta-ualr/NeuWalk. The version used in this study is archived on Zenodo at https://doi.org/10.5281/zenodo.22929014 and released under the Creative Commons Attribution 4.0 International (CC BY 4.0) license. All experimental reconstructions analyzed in this study are publicly available on NeuroMorpho.Org [38]. Supplementary material accompanying this article can be found in the online version of the manuscript.

## Declaration of generative AI and AI-assisted technologies in the manuscript preparation process

During the preparation of this work, the author used Claude (Anthropic) in order to assist with manuscript editing, including grammar and consistency checking, bibliography verification, table formatting, and content review. Additionally, the same tool assisted with the development of the NeuWalk package. The author reviewed and edited the content as well as the code as needed, and takes full responsibility for the content of the published article and for the associated source code.

---

**Algorithm 1.**
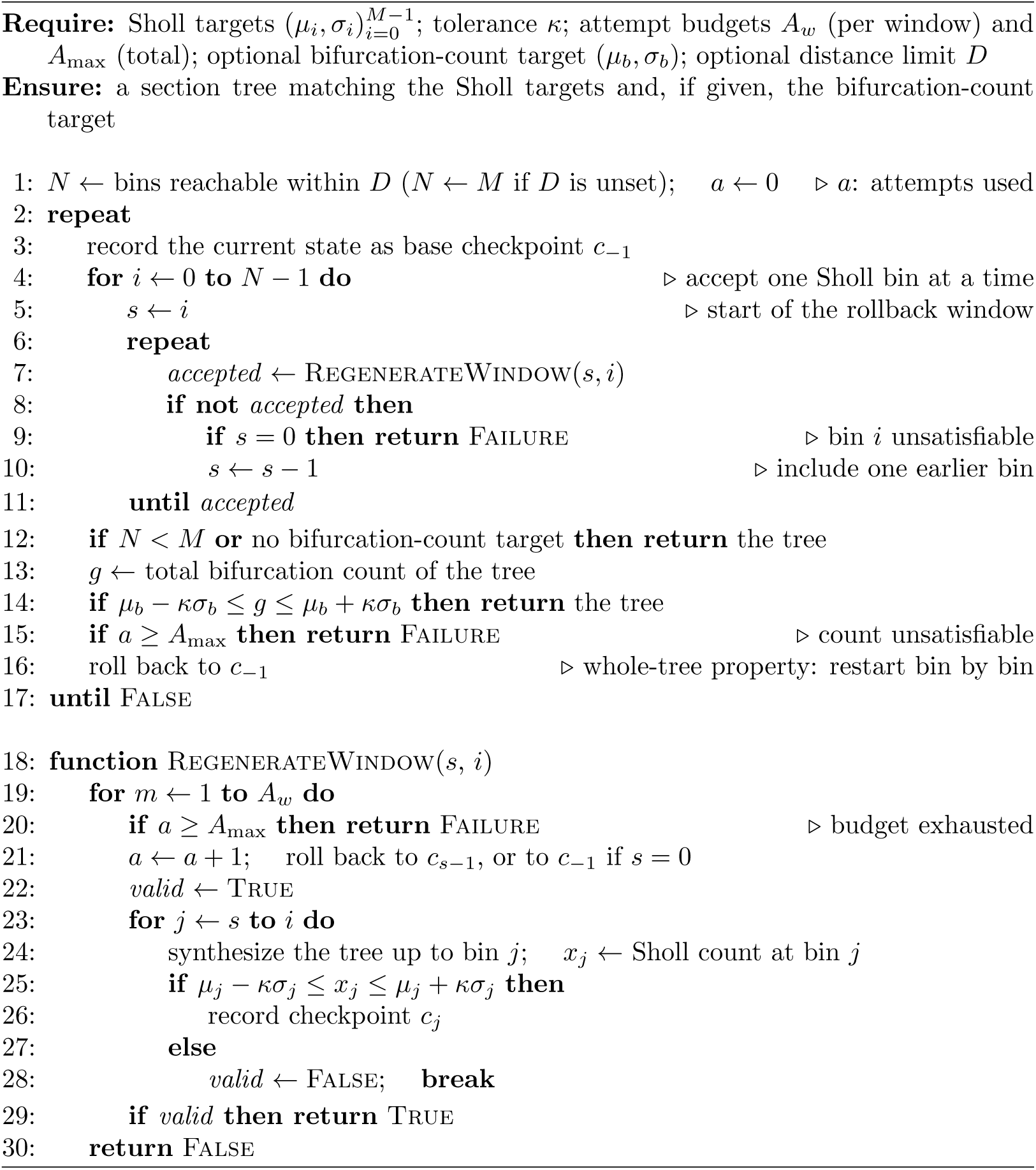
Progressive topology synthesis.

**Algorithm 2.**
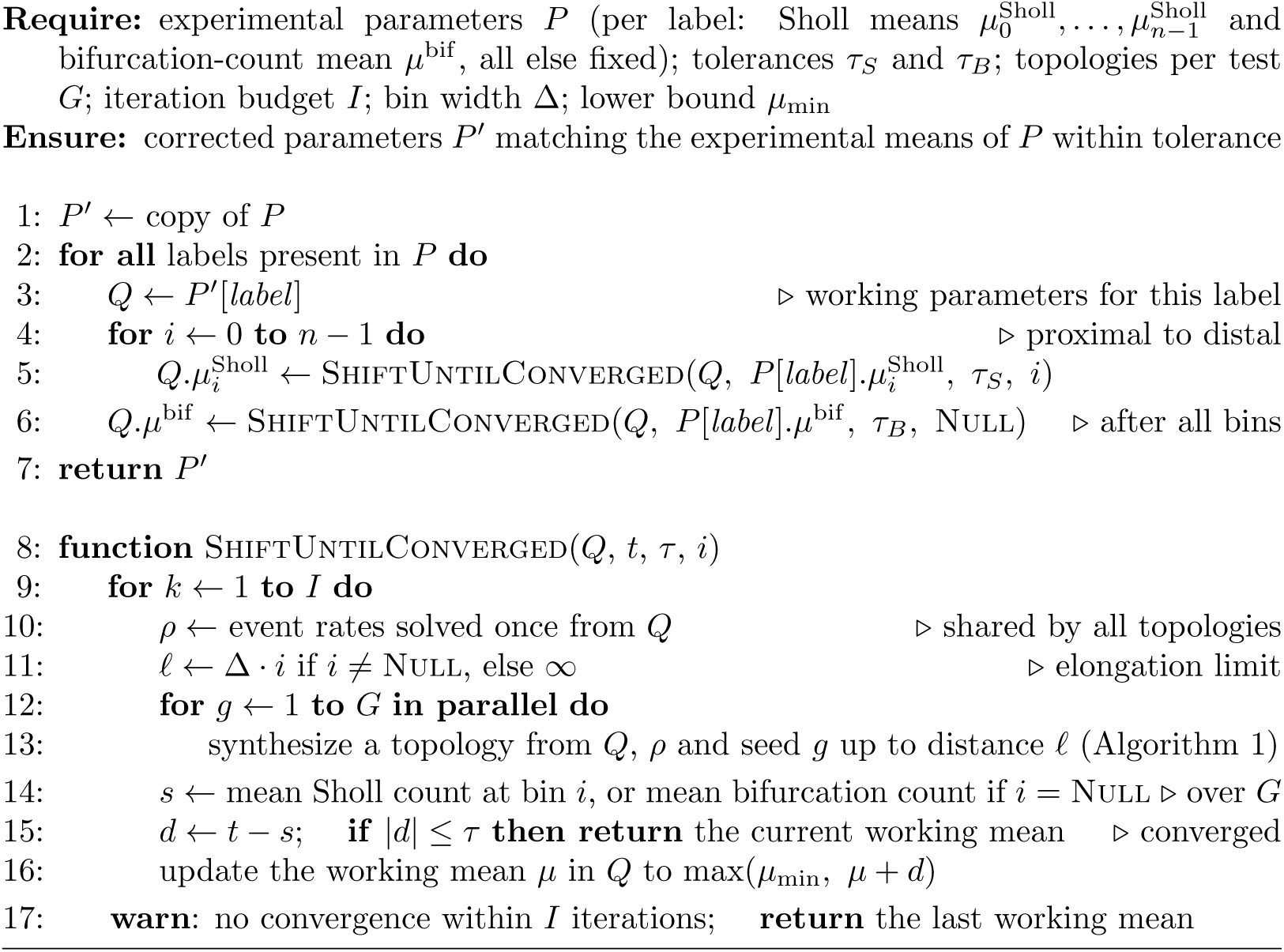
Two-stage calibration of synthesis parameters.

## Notes

http://www.neuromorpho.org

### Competing Interest Statement

The authors have declared no competing interest.

