## Supplementary Material for "NeuWalk: A Python Library for Data-Driven Synthesis of Neuron Morphologies Using Random Walks"

NeuWalk: A Python Package for Data-Driven Synthesis of Three-Dimensional Neuron Morphologies Using Random Walks

Francesco Cavarretta

Department of Computer Science and Emerging Analytics Center, University of Arkansas at Little Rock, Little Rock, AR, United States

#### S1 Morphology curation

We curated semilunar (SL) and pyramidal (PYR) cells from the anterior piriform cortex. For SLs, only the apical dendritic tree was analyzed. In one cell, we corrected apical dendrites that appeared mislabeled (d080305a-100). For PYRs, four morphologies were discarded from the original dataset because they exhibited severe deformation or amputation in their dendrites, leaving twelve morphologies for our analysis.

In four MC reconstructions (2M3, TM2, TM3, and TM5), we corrected mislabeled basal dendrites and reconnected them directly to the soma. Similarly, the apical dendrite and tuft were incorrectly labeled as basal dendrites across all mTC reconstructions and were relabeled accordingly. In two mTC reconstructions (2T2 and TT3), we noted that a single short basal dendritic segment emerged directly from the soma (7.4 and 12.2 $\mu$m in length, respectively). As these segments were likely artifacts, they were removed, and their child branches were reconnected directly to the soma.

The Markram dataset [1] comprises several hundred morphological reconstructions from rat somatosensory cortex, including 164 layer 5 pyramidal neurons. Layer 5 pyramidal neurons are further divided into several subtypes, distinguished primarily by the size and branching complexity of the apical tuft: thick-tufted pyramidal cells (TTPCs), bearing a prominent apical dendrite terminating in a wide, elaborately branched tuft and projecting primarily to subcortical targets; slender-tufted pyramidal cells (STPCs), with a comparatively smaller and less branched tuft; and untufted pyramidal cells (UTPCs), lacking a distinct apical tuft altogether [2, 1]. Among these, TTPCs are widely regarded as the most morphologically complex and extensively studied layer 5 subtype [2]. We therefore focused our testing on TTPCs, selecting exemplar reconstructions from the archive in order to probe the method under the most morphologically demanding conditions available. Because curation was performed manually, and because a small number of reconstructions proved sufficient to estimate parameter values reliably, we selected $n = 10$ TTPC reconstructions.

### S2 Biased-and-correlated random walk

#### S2.1 Elongation

At each elongation, the position of the new tip $p_{i+1}$ is computed as

$$p_{i+1} = p_i + \lambda \vec{v}_{i+1}$$

where $p_i$ is the current tip position, $\vec{v}_{i+1}$ is the updated unit direction vector, and $\lambda$ is the step length, corrected at each iteration so that the tip advances a fixed distance $\Lambda$.

Given a neurite, $\vec{c}_i$ describes its state at the $i$-th iteration. It encodes multiple descriptors, whose full details are not necessary at this stage. In particular, it comprises $\vec{v}_i$ and $p_i$, as defined above. Let $\mathcal{C}$ denote the set of admissible neurite states. A bias function is defined as a map

$$\mathcal{B} : \mathcal{C} \to \mathbb{R}^3$$

which takes the current neurite state $\vec{c} \in \mathcal{C}$ and returns a three-dimensional bias vector

$$\vec{b} = \mathcal{B}(\vec{c})$$

The vector $\vec{b}$ represents the directional contribution induced by the corresponding bias.

More sophisticated bias functions can be derived by combining simpler ones. Given a family of bias functions $\mathcal{B}_1, \ldots, \mathcal{B}_M$ and weights $w^g = (w_1^g, \ldots, w_M^g)$, we define a composite bias function by the linear combination

$$\mathcal{B}_{w^g}(\vec{c}) = \sum_{k=1}^{M} w_k^g \mathcal{B}_k(\vec{c})$$

We now introduce a mathematical operator $\otimes$, referred to here as the sequential combination operator. This operator adds a vector to a unit vector and projects the result back onto the unit sphere. Let

$$\mathbb{S}^2 = \left\{ \vec{x} \in \mathbb{R}^3 : \|\vec{x}\| = 1 \right\}$$

denote the unit sphere in three-dimensional space. We define

$$\otimes : \mathbb{S}^2 \times \mathbb{R}^3 \to \mathbb{S}^2$$

as

$$\vec{x} \otimes \vec{y} = \frac{\vec{x} + \vec{y}}{\|\vec{x} + \vec{y}\|}$$

provided that $\vec{x} + \vec{y} \neq \vec{0}$. Note that the sequential application of this operator is order-dependent. Also, if $\|\vec{y}\| \gg \|\vec{x}\|$, then

$$\vec{x} \otimes \vec{y} \to \frac{\vec{y}}{\|\vec{y}\|}$$

Thus, the norms of the vectors must be calibrated carefully to avoid one dominating the others. We will refer to this as the *degeneration property*.

Given a family of bias vectors $\{\vec{b}_j\}_{j=1}^N$ with weights $\{w_j^b\}_{j=1}^N$, and a random vector $\vec{r}$ with weight $w^r$, the updated direction $\vec{v}_{i+1}$ is computed as

$$\vec{v}_{i+1} = \vec{v}_i \otimes w_1^b \vec{b}_1 \otimes w_2^b \vec{b}_2 \cdots \otimes w_N^b \vec{b}_N \otimes w^r \vec{r}$$

where

$$\vec{b}_j = \mathcal{B}_j(\vec{c}_i), \qquad j = 1, \ldots, N$$

#### S2.2 Bifurcation bias

Bifurcation events require a distinct mechanism from elongation, since they must produce two new initial directions — one for each offspring — rather than update a single ongoing direction. We define a bifurcation bias as a map

$$\mathcal{D} : \mathcal{C} \to \mathbb{S}^2 \times \mathbb{S}^2$$

taking the parent tip's state at the moment of branching and returning a pair of initial directions $(\vec{v}_1^{(\text{left})}, \vec{v}_1^{(\text{right})})$ for the two offspring. Unlike elongation biases, bifurcation biases are not composed via $\otimes$ or linear combination; a single bifurcation bias determines both offspring directions directly from the parent's incoming direction.

Given the last direction $\vec{v}_{\text{last}}^{(\text{parent})}$ of the torsion force $\vec{R}$, a lateral direction $\hat{\ell} = \left(\vec{v}_i \times \vec{R}\right) / \|\vec{v}_i \times \vec{R}\|$ is obtained, orthogonal to $\vec{v}_{\text{last}}^{(\text{parent})}$, and the offspring directions follow as

$$\vec{v}_0^{(\text{left})} = \cos\theta^{(\text{left})}\, \vec{v}_{\text{last}}^{(\text{parent})} - \sin\theta^{(\text{left})}\, \hat{\ell}$$

and

$$\vec{v}_0^{(\text{right})} = \cos\theta^{(\text{right})}\, \vec{v}_{\text{last}}^{(\text{parent})} + \sin\theta^{(\text{right})}\, \hat{\ell}$$

where $\theta^{(\text{left})}$ and $\theta^{(\text{right})}$ are the bifurcation angles of the two offspring, and the normal to the bifurcation plane is proportional to the component of $\vec{R}$ orthogonal to $\vec{v}_{\text{last}}^{(\text{parent})}$. In `radial_torsion`, $\vec{u}$ is the aggregate repulsion exerted by neighboring dendrites and $\theta^{(\text{left})} = \theta^{(\text{right})}$; in `cross_torsion`, $\vec{R}$ is the surface normal of an ellipsoid at the bifurcation point; in `internal_branch`, $\theta^{(\text{left})} = 0$, so that one offspring continues along the parent direction, and $\hat{\ell}$ is drawn with a random azimuth.

#### S3 Sholl intersections by bin

Tables S1–S5 report the bin-by-bin comparison of Sholl intersections between natural and synthetic populations, for each neuron class and dendritic arbor analyzed in the main text.

Table S1: Sholl intersections by bin for middle tufted cells (mTCs), natural (Nat) vs. synthetic (Synth) populations, reported as mean ± SD. Bins are 50 $\mu$m wide. $p_{\text{mean}}$ and $p_{\text{var}}$ are two-tailed bootstrap $p$-values (10,000 resamples) for the difference in mean and in variance, respectively. Asterisks denote differences that were both significant after Bonferroni correction across the bins tested ($\alpha = 0.01/19 = 0.0005$ for the basal arbor) and above the minimum effect size defined in the main text. Bins in which one population had zero variance were not tested, as the variance ratio is undefined and the resampling distribution of the mean is degenerate; both $p$-values are reported as — for these bins. Bins reached by fewer than three neurons per population are omitted.

| Bin ($\mu$m) | Nat | Synth | $p_{\text{mean}}$ | $p_{\text{var}}$ |
|---|---|---|---|---|
| 0–50 | 3.80 ± 1.60 | 4.20 ± 1.47 | 0.588 | 0.365 |
| 50–100 | 6.20 ± 2.32 | 7.14 ± 2.66 | 0.392 | 0.862 |
| 100–150 | 8.00 ± 3.46 | 8.40 ± 2.73 | 0.802 | 0.185 |
| 150–200 | 7.80 ± 2.56 | 8.72 ± 2.65 | 0.458 | 0.714 |
| 200–250 | 9.00 ± 1.26 | 9.71 ± 2.16 | 0.233 | 0.198 |
| 250–300 | 8.80 ± 2.64 | 9.21 ± 2.15 | 0.750 | 0.331 |
| 300–350 | 9.20 ± 1.94 | 9.53 ± 2.30 | 0.719 | 0.823 |
| 350–400 | 8.60 ± 1.96 | 9.43 ± 2.35 | 0.358 | 0.769 |
| 400–450 | 8.80 ± 1.72 | 9.53 ± 2.40 | 0.376 | 0.413 |
| 450–500 | 8.40 ± 1.50 | 9.12 ± 2.25 | 0.308 | 0.356 |
| 500–550 | 8.00 ± 1.10 | 9.01 ± 1.91 | 0.040 | 0.327 |
| 550–600 | 9.40 ± 1.50 | 9.23 ± 2.01 | 0.793 | 0.504 |
| 600–650 | 8.00 ± 1.67 | 8.70 ± 2.03 | 0.366 | 0.732 |
| 650–700 | 5.60 ± 3.44 | 6.50 ± 1.98 | 0.572 | 0.308 |
| 700–750 | 4.00 ± 1.90 | 5.60 ± 1.95 | 0.062 | 0.685 |
| 750–800 | 2.80 ± 1.17 | 4.11 ± 1.75 | 0.010 | 0.311 |
| 800–850 | 1.40 ± 1.02 | 1.63 ± 1.32 | 0.627 | 0.622 |
| 850–900 | 0.80 ± 0.40 | 0.32 ± 0.66 | 0.007 | 0.339 |
| 900–950 | 0.80 ± 0.40 | 0.06 ± 0.35 | 0.001 | 0.634 |

Table S2: Sholl intersections by bin for mitral cells (MCs), natural (Nat) vs. synthetic (Synth) populations, reported as mean ± SD. Bins are 50 $\mu$m wide. $p_{\text{mean}}$ and $p_{\text{var}}$ are two-tailed bootstrap $p$-values (10,000 resamples) for the difference in mean and in variance, respectively. Asterisks denote differences that were both significant after Bonferroni correction across the bins tested ($\alpha = 0.01/22 = 0.0005$ for the basal arbor) and above the minimum effect size defined in the main text. Bins in which one population had zero variance were not tested, as the variance ratio is undefined and the resampling distribution of the mean is degenerate; both $p$-values are reported as — for these bins. Bins reached by fewer than three neurons per population are omitted.

| Bin ($\mu$m) | Nat | Synth | $p_{\text{mean}}$ | $p_{\text{var}}$ |
|---|---|---|---|---|
| 0–50 | 6.22 ± 1.23 | 6.51 ± 1.34 | 0.514 | 0.853 |
| 50–100 | 7.89 ± 0.87 | 9.04 ± 2.48 | 0.001 | 0.029 |
| 100–150 | 10.67 ± 2.05 | 10.12 ± 2.76 | 0.447 | 0.202 |
| 150–200 | 13.00 ± 2.40 | 12.49 ± 3.16 | 0.550 | 0.197 |
| 200–250 | 13.33 ± 2.05 | 13.54 ± 3.30 | 0.778 | 0.176 |
| 250–300 | 14.33 ± 2.67 | 14.12 ± 3.35 | 0.822 | 0.515 |
| 300–350 | 15.67 ± 2.71 | 15.42 ± 3.59 | 0.790 | 0.419 |
| 350–400 | 15.67 ± 3.16 | 15.63 ± 3.58 | 0.969 | 0.705 |
| 400–450 | 15.78 ± 3.49 | 16.21 ± 3.76 | 0.719 | 0.915 |
| 450–500 | 15.56 ± 4.11 | 16.18 ± 3.67 | 0.657 | 0.289 |
| 500–550 | 15.22 ± 3.82 | 15.43 ± 3.70 | 0.882 | 0.671 |
| 550–600 | 13.67 ± 3.83 | 14.21 ± 3.57 | 0.674 | 0.488 |
| 600–650 | 12.33 ± 3.37 | 12.71 ± 3.44 | 0.753 | 0.859 |
| 650–700 | 11.78 ± 2.74 | 11.46 ± 3.20 | 0.737 | 0.538 |
| 700–750 | 9.89 ± 2.69 | 11.16 ± 3.07 | 0.173 | 0.681 |
| 750–800 | 8.89 ± 3.07 | 9.16 ± 2.85 | 0.802 | 0.432 |
| 800–850 | 7.11 ± 3.21 | 7.36 ± 2.53 | 0.824 | 0.306 |
| 850–900 | 5.00 ± 2.58 | 5.42 ± 2.29 | 0.635 | 0.189 |
| 900–950 | 3.22 ± 1.93 | 3.46 ± 2.14 | 0.726 | 0.815 |
| 950–1000 | 2.44 ± 1.34 | 2.42 ± 1.71 | 0.959 | 0.468 |
| 1000–1050 | 1.22 ± 0.92 | 1.20 ± 1.57 | 0.936 | 0.088 |
| 1050–1100 | 0.89 ± 1.10 | 0.22 ± 0.70 | 0.066 | 0.075 |

Table S3: Sholl intersections by bin for semilunar cells (SLs), natural (Nat) vs. synthetic (Synth) populations, reported as mean ± SD. Bins are 50 $\mu$m wide. $p_{\text{mean}}$ and $p_{\text{var}}$ are two-tailed bootstrap $p$-values (10,000 resamples) for the difference in mean and in variance, respectively. Asterisks denote differences that were both significant after Bonferroni correction across the bins tested ($\alpha = 0.01/8 = 0.001$ for the apical arbor) and above the minimum effect size defined in the main text. Bins in which one population had zero variance were not tested, as the variance ratio is undefined and the resampling distribution of the mean is degenerate; both $p$-values are reported as — for these bins. Bins reached by fewer than three neurons per population are omitted.

| Bin ($\mu$m) | Nat | Synth | $p_{\text{mean}}$ | $p_{\text{var}}$ |
|---|---|---|---|---|
| 0–50 | 2.33 ± 0.47 | 2.37 ± 0.48 | 0.670 | 0.332 |
| 50–100 | 6.00 ± 1.41 | 5.56 ± 2.11 | 0.560 | 0.333 |
| 100–150 | 8.00 ± 0.82 | 7.86 ± 1.63 | 0.777 | 0.548 |
| 150–200 | 9.67 ± 4.11 | 9.72 ± 2.20 | 0.949 | 0.136 |
| 200–250 | 10.33 ± 4.03 | 10.52 ± 2.52 | 0.787 | 0.333 |
| 250–300 | 11.33 ± 4.03 | 11.51 ± 2.60 | 0.806 | 0.326 |
| 300–350 | 10.00 ± 0.82 | 10.08 ± 1.53 | 0.864 | 0.553 |
| 350–400 | 6.67 ± 2.49 | 6.94 ± 1.56 | 0.783 | 0.334 |

Table S4: Sholl intersections by bin for pyramidal neurons (PYRs) of the anterior piriform cortex, natural (Nat) vs. synthetic (Synth) populations, reported as mean ± SD. Bins are 50 $\mu$m wide. $p_{\text{mean}}$ and $p_{\text{var}}$ are two-tailed bootstrap $p$-values (10,000 resamples) for the difference in mean and in variance, respectively. Asterisks denote differences that were both significant after Bonferroni correction across the bins tested ($\alpha = 0.01/13 = 0.0008$ for the apical arbor, $\alpha = 0.01/10 = 0.001$ for the basal arbor) and above the minimum effect size defined in the main text. Bins in which one population had zero variance were not tested, as the variance ratio is undefined and the resampling distribution of the mean is degenerate; both $p$-values are reported as — for these bins. Bins reached by fewer than three neurons per population are omitted.

| Bin ($\mu$m) | Nat | Synth | $p_{\text{mean}}$ | $p_{\text{var}}$ |
|---|---|---|---|---|
| *Apical dendrites* | | | | |
| 0–50 | 1.08 ± 0.28 | 1.00 ± 0.00 | — | — |
| 50–100 | 2.92 ± 1.38 | 3.09 ± 1.53 | 0.685 | 0.770 |
| 100–150 | 4.00 ± 1.96 | 4.36 ± 2.11 | 0.546 | 0.863 |
| 150–200 | 5.17 ± 2.34 | 5.62 ± 2.68 | 0.521 | 0.589 |
| 200–250 | 6.75 ± 2.89 | 8.28 ± 3.65 | 0.074 | 0.572 |
| 250–300 | 8.17 ± 3.13 | 10.32 ± 4.51 | 0.023 | 0.145 |
| 300–350 | 9.25 ± 3.29 | 11.57 ± 4.94 | 0.022 | 0.067 |
| 350–400 | 10.33 ± 3.86 | 12.93 ± 5.41 | 0.029 | 0.324 |
| 400–450 | 8.08 ± 3.88 | 10.08 ± 4.36 | 0.085 | 0.645 |
| 450–500 | 7.58 ± 5.12 | 9.56 ± 4.26 | 0.191 | 0.162 |
| 500–550 | 7.08 ± 5.17 | 8.88 ± 3.98 | 0.252 | 0.063 |
| 550–600 | 6.17 ± 6.05 | 7.59 ± 3.45 | 0.435 | 0.012 |
| 600–650 | 3.25 ± 3.51 | 2.94 ± 2.44 | 0.764 | 0.047 |
| 650–700 | 0.67 ± 1.03 | 0.55 ± 0.77 | 0.701 | 0.158 |
| *Basal dendrites* | | | | |
| 0–50 | 5.25 ± 1.88 | 5.55 ± 1.81 | 0.604 | 0.506 |
| 50–100 | 10.25 ± 5.55 | 10.40 ± 4.47 | 0.931 | 0.515 |
| 100–150 | 12.50 ± 4.35 | 12.90 ± 5.62 | 0.776 | 0.294 |
| 150–200 | 11.75 ± 3.85 | 12.21 ± 5.46 | 0.703 | 0.120 |
| 200–250 | 8.83 ± 2.88 | 9.09 ± 4.19 | 0.777 | 0.087 |
| 250–300 | 5.92 ± 2.75 | 5.92 ± 3.03 | 0.997 | 0.722 |
| 300–350 | 4.00 ± 3.14 | 3.85 ± 2.49 | 0.872 | 0.236 |
| 350–400 | 2.25 ± 2.59 | 2.02 ± 2.11 | 0.766 | 0.276 |
| 400–450 | 1.33 ± 1.49 | 0.98 ± 1.54 | 0.433 | 0.943 |
| 450–500 | 0.75 ± 1.16 | 0.12 ± 0.51 | 0.058 | 0.033 |

Table S5: Sholl intersections by bin for neocortical pyramidal neurons (NCPYRs), natural (Nat) vs. synthetic (Synth) populations, reported as mean ± SD. Bins are 50 $\mu$m wide. $p_{\text{mean}}$ and $p_{\text{var}}$ are two-tailed bootstrap $p$-values (10,000 resamples) for the difference in mean and in variance, respectively. Asterisks denote differences that were both significant after Bonferroni correction across the bins tested ($\alpha = 0.01/19 = 0.0005$ for the apical + oblique arbor, $\alpha = 0.01/5 = 0.002$ for the basal arbor) and above the minimum effect size defined in the main text. Bins in which one population had zero variance were not tested, as the variance ratio is undefined and the resampling distribution of the mean is degenerate; both $p$-values are reported as — for these bins. Bins reached by fewer than three neurons per population are omitted.

| Bin ($\mu$m) | Nat | Synth | $p_{\text{mean}}$ | $p_{\text{var}}$ |
|---|---|---|---|---|
| *Apical + oblique dendrites* | | | | |
| 0–50 | 1.00 ± 0.00 | 1.00 ± 0.00 | — | — |
| 50–100 | 6.60 ± 2.24 | 5.12 ± 3.04 | 0.044 | 0.161 |
| 100–150 | 11.40 ± 4.10 | 8.87 ± 5.00 | 0.057 | 0.530 |
| 150–200 | 11.00 ± 4.45 | 10.43 ± 4.78 | 0.697 | 0.917 |
| 200–250 | 9.20 ± 2.60 | 8.63 ± 4.47 | 0.521 | 0.033 |
| 250–300 | 4.00 ± 2.72 | 5.36 ± 3.22 | 0.127 | 0.566 |
| 300–350 | 1.80 ± 1.78 | 2.67 ± 1.96 | 0.104 | 0.827 |
| 350–400 | 1.80 ± 0.98 | 1.64 ± 0.95 | 0.627 | 0.771 |
| 400–450 | 1.90 ± 0.70 | 1.66 ± 0.80 | 0.293 | 0.640 |
| 450–500 | 2.10 ± 0.54 | 1.85 ± 0.84 | 0.168 | 0.156 |
| 500–550 | 2.50 ± 0.67 | 2.12 ± 1.04 | 0.091 | 0.144 |
| 550–600 | 2.70 ± 0.90 | 2.28 ± 1.18 | 0.147 | 0.500 |
| 600–650 | 3.20 ± 0.75 | 2.67 ± 1.35 | 0.036 | 0.175 |
| 650–700 | 4.00 ± 1.00 | 3.36 ± 1.79 | 0.065 | 0.036 |
| 700–750 | 4.50 ± 1.12 | 3.87 ± 1.76 | 0.096 | 0.036 |
| 750–800 | 5.90 ± 2.66 | 6.02 ± 2.95 | 0.896 | 0.815 |
| 800–850 | 8.50 ± 3.61 | 8.68 ± 4.21 | 0.885 | 0.668 |
| 850–900 | 9.60 ± 3.17 | 9.89 ± 4.72 | 0.792 | 0.129 |
| 900–950 | 10.90 ± 6.53 | 11.38 ± 5.57 | 0.824 | 0.273 |
| 950–1000 | 8.50 ± 10.03 | 10.20 ± 5.65 | 0.599 | 0.035 |
| *Basal dendrites* | | | | |
| 0–50 | 7.80 ± 1.40 | 8.02 ± 1.57 | 0.642 | 0.769 |
| 50–100 | 24.50 ± 8.92 | 22.39 ± 9.84 | 0.478 | 0.804 |
| 100–150 | 24.50 ± 10.76 | 24.15 ± 10.81 | 0.920 | 0.809 |
| 150–200 | 14.00 ± 7.82 | 13.71 ± 6.42 | 0.901 | 0.278 |
| 200–250 | 2.50 ± 1.80 | 2.53 ± 2.13 | 0.962 | 0.595 |

#### S4 NCPYR validation by dendritic subclass

Because the NCPYR apical dendrite is synthesized using the subclass-specific trunk/oblique strategy described in the main text, we additionally validated Sholl intersections, bifurcation count, and total dendritic length separately for the apical trunk and the apical oblique dendrites. Tables S6 and S7 report these comparisons, and Table S8 reports the corresponding bin-by-bin Sholl comparison; the corresponding tables in the main text, and Table S5, instead report the apical arbor as a whole (trunk and oblique combined).

Table S6: Bifurcation count for neocortical pyramidal neurons (NCPYR), natural (Nat) vs. synthetic (Synth) populations, reported as mean ± SD, broken down by apical dendritic subclass. $n = 10$ for both subclasses. $p_{\text{mean}}$ and $p_{\text{var}}$ are two-tailed bootstrap $p$-values (10,000 resamples, $\alpha = 0.01$) for the difference in mean and in variance, respectively.

| Subclass | Nat | Synth | $p_{\text{mean}}$ | $p_{\text{var}}$ |
|---|---|---|---|---|
| Apical (trunk) | 25.5 ± 8.7 | 26.4 ± 14.0 | 0.772 | 0.273 |
| Apical oblique | 17.3 ± 7.0 | 16.9 ± 7.2 | 0.865 | 0.925 |

Table S7: Total dendritic length ($\mu$m) for neocortical pyramidal neurons (NCPYR), natural (Nat) vs. synthetic (Synth) populations, reported as mean ± SD, broken down by apical dendritic subclass. $n = 10$ for both subclasses. $p_{\text{mean}}$ and $p_{\text{var}}$ are two-tailed bootstrap $p$-values (10,000 resamples, $\alpha = 0.01$) for the difference in mean and in variance, respectively.

| Subclass | Nat | Synth | $p_{\text{mean}}$ | $p_{\text{var}}$ |
|---|---|---|---|---|
| Apical (trunk) | 5113.4 ± 1215.0 | 4917.4 ± 1838.1 | 0.629 | 0.120 |
| Apical oblique | 3196.9 ± 723.4 | 3136.0 ± 997.6 | 0.803 | 0.150 |

Table S8: Sholl intersections by bin for neocortical pyramidal neurons (NCPYRs), with the apical trunk and the oblique dendrites analyzed separately, natural (Nat) vs. synthetic (Synth) populations, reported as mean ± SD. Bins are 50 $\mu$m wide. $p_{\text{mean}}$ and $p_{\text{var}}$ are two-tailed bootstrap $p$-values (10,000 resamples) for the difference in mean and in variance, respectively. Asterisks denote differences that were both significant after Bonferroni correction across the bins tested ($\alpha = 0.01/14 = 0.0007$ for the apical trunk arbor, $\alpha = 0.01/5 = 0.002$ for the apical oblique arbor) and above the minimum effect size defined in the main text. Bins in which one population had zero variance were not tested, as the variance ratio is undefined and the resampling distribution of the mean is degenerate; both $p$-values are reported as — for these bins. Bins reached by fewer than three neurons per population are omitted. For the oblique dendrites, distances are measured from their point of origin on the apical trunk.

| Bin ($\mu$m) | Nat | Synth | $p_{\text{mean}}$ | $p_{\text{var}}$ |
|---|---|---|---|---|
| *Apical trunk dendrites* | | | | |
| 0–50 | 1.00 ± 0.00 | 1.00 ± 0.00 | — | — |
| 50–100 | 1.00 ± 0.00 | 1.00 ± 0.00 | — | — |
| 100–150 | 1.00 ± 0.00 | 1.00 ± 0.00 | — | — |
| 150–200 | 1.00 ± 0.00 | 1.00 ± 0.00 | — | — |
| 200–250 | 1.00 ± 0.00 | 1.00 ± 0.00 | — | — |
| 250–300 | 1.00 ± 0.00 | 1.00 ± 0.00 | — | — |
| 300–350 | 1.10 ± 0.30 | 1.10 ± 0.29 | 0.925 | 0.787 |
| 350–400 | 1.30 ± 0.46 | 1.23 ± 0.42 | 0.670 | 0.318 |
| 400–450 | 1.80 ± 0.60 | 1.60 ± 0.73 | 0.293 | 0.561 |
| 450–500 | 2.10 ± 0.54 | 1.84 ± 0.83 | 0.152 | 0.157 |
| 500–550 | 2.50 ± 0.67 | 2.12 ± 1.04 | 0.091 | 0.144 |
| 550–600 | 2.70 ± 0.90 | 2.28 ± 1.18 | 0.147 | 0.500 |
| 600–650 | 3.20 ± 0.75 | 2.67 ± 1.35 | 0.036 | 0.175 |
| 650–700 | 4.00 ± 1.00 | 3.36 ± 1.79 | 0.065 | 0.036 |
| 700–750 | 4.50 ± 1.12 | 3.87 ± 1.76 | 0.096 | 0.036 |
| 750–800 | 5.90 ± 2.66 | 6.02 ± 2.95 | 0.896 | 0.815 |
| 800–850 | 8.50 ± 3.61 | 8.68 ± 4.21 | 0.885 | 0.668 |
| 850–900 | 9.60 ± 3.17 | 9.89 ± 4.72 | 0.792 | 0.129 |
| 900–950 | 10.90 ± 6.53 | 11.38 ± 5.57 | 0.824 | 0.273 |
| 950–1000 | 8.50 ± 10.03 | 10.20 ± 5.65 | 0.599 | 0.035 |
| *Apical oblique dendrites* | | | | |
| 0–50 | 10.50 ± 1.96 | 10.77 ± 2.00 | 0.684 | 0.879 |
| 50–100 | 20.90 ± 5.56 | 20.37 ± 6.91 | 0.770 | 0.200 |
| 100–150 | 15.00 ± 4.47 | 15.12 ± 5.36 | 0.940 | 0.552 |
| 150–200 | 5.40 ± 2.84 | 5.58 ± 2.87 | 0.853 | 0.854 |
| 200–250 | 0.70 ± 0.90 | 0.60 ± 0.73 | 0.748 | 0.142 |
